# The Origin Recognition Complex requires chromatin tethering by a hypervariable intrinsically disordered region that is functionally conserved from sponge to man

**DOI:** 10.1101/2023.05.11.540405

**Authors:** Olubu A. Adiji, Brendan S. McConnell, Matthew W. Parker

## Abstract

The first step towards eukaryotic genome duplication is loading of the replicative helicase, the Mcm2-7 complex, onto chromatin. This so-called “licensing” step is executed by the Pre-Replication Complex (Pre-RC) whose assembly on chromatin is nucleated by the DNA-binding activity of the Origin Recognition Complex (ORC). It is thought that metazoan ORC, like the yeast complex, is recruited directly to chromatin by its ATP-dependent DNA binding and encirclement activity. However, we have previously shown that this DNA binding mode is dispensable for chromatin recruitment of fly ORC, raising the question of how metazoan ORC binds chromosomes. We show here that the intrinsically disordered region (IDR) of fly Orc1 is both necessary and sufficient for recruitment of ORC to chromosomes *in vivo* and demonstrate that this activity is regulated by IDR phosphorylation. *In vitro* studies show that the IDR alone binds DNA and this bestows the ORC holocomplex with a high-affinity *ATP-independent* DNA binding mode. Interestingly, we find that Orc1 IDRs have diverged so markedly across metazoans that they are unrecognizable as orthologs and yet we find that these compositionally homologous sequences retain DNA and chromatin binding activity down to basal metazoans. Altogether, these data suggest that chromatin is recalcitrant to ORC’s ATP-dependent DNA binding activity and we propose that this necessitates IDR-dependent chromatin tethering which poises ORC to opportunistically encircle nucleosome free regions as they become available. This work reveals a novel step in metazoan replication licensing and expands our understanding of disordered protein homology and evolution by stretching the relationship between primary structure and function.

## INTRODUCTION

Eukaryotic DNA replication initiation occurs in two temporally separate steps known as licensing and firing. Replication licensing constitutes loading of the replicative helicase, the heterohexameric Mcm2-7 complex, onto replication start sites, or “origins”, during late mitosis and early G1 phase of the cell cycle (reviewed in (1)). During the Synthesis (S) phase of the cell cycle, Mcm2-7 scaffolds replisome assembly and replication initiates, or “fires”, leading to DNA strand separation and duplication of cellular genetic material. *In vitro* reconstitution studies have led to a precise mechanistic understanding of the DNA replication licensing reaction. In the first step, the Origin Recognition Complex (ORC, composed of Orc1-6) binds and encircles DNA. This nucleates assembly of the Pre-Replication Complex (Pre-RC), a macromolecular machine composed of ORC, Cdc6, Cdt1 and Mcm2-7. The Pre-RC then loads Mcm2-7 around duplex DNA as a double hexamer (2,3) where it is poised to be activated in S-phase.

The first step of origin licensing is recruitment of ORC to chromatin. This process has been studied intensely in budding yeast (*S. cerevisiae*) and with recombinant fly and human ORC. A common theme across species is the formation of an ATP-dependent ORC•DNA ternary complex (4–8) which structural studies show manifests as DNA encirclement within the ORC central channel (8–10). In addition to being an essential intermediate in Pre-RC assembly, this DNA binding mode is also thought to mediate the initial recruitment of ORC to chromatin. This mechanism of DNA binding is consistent with yeast ORC’s constitutive association with chromosomes (11) but it is less obvious how it relates to the regulated chromatin association observed for metazoan ORC (12,13). This mechanism is contrasted with that of *S. pombe* ORC, which utilizes a two-step mechanism of chromatin binding. In the first step, *Sp*ORC is tethered to chromatin via *ATP-independent* DNA binding facilitated by AT-hook motifs embedded within the *Sp*Orc4 N-terminus (14–17). In the second step, the assembly transitions to a salt-stable complex that, based on conservation of the ORC core complex, likely represents DNA encirclement (18). Why *Sp*ORC requires a two-step mechanism, while *S. cerevisiae* and metazoan ORC appear to proceed by direct ATP-dependent DNA binding and encirclement, is currently unclear.

We previously observed that *D. melanogaster* ORC’s ATP-binding (Walker A) and hydrolysis motifs (Walker B) – regions necessary for ATP-dependent DNA binding *in vitro* (5) – are dispensable for the recruitment of ORC to chromosomes *in vivo* (19). Consistently, a large fraction of recombinant fly and human ORC’s *in vitro* DNA binding activity is ATP-independent (5,7,20). These data suggest that metazoan ORC may first be tethered to chromatin via an *ATP-independent* mechanism, reminiscent of the two-step binding path of *S. pombe* ORC. Indeed, metazoan ORC contains multiple DNA and chromatin binding elements that could, in theory, facilitate this. For example, we previously identified an intrinsically disordered region (IDR) in metazoan Orc1 that binds and phase separates with DNA *in vitro* (19). N-terminal to the Orc1 IDR is a Bromo Adjacent Homology (BAH) domain that in chordates binds histone H4 dimethylated at lysine 20 (21) and is important for chromatin localization (22). Finally, the TFIIB-like domains of *D. melanogaster* Orc6 bind DNA *in vitro* (23). It is currently unknown which of these chromatin binding elements, if any, are required for chromatin tethering of metazoan ORC.

Here we demonstrate that the fly Orc1 IDR is necessary and sufficient for recruitment of ORC to chromosomes *in vivo*. We show that the Orc1 IDR can directly bind both DNA and chromatin and that Cyclin-dependent kinases (CDKs) regulate these interactions through IDR phosphorylation. Combined with our previous work, these data strongly suggest that chromatin is recalcitrant to ATP-dependent DNA encirclement by ORC but not IDR-dependent chromatin recruitment. Interestingly, we find that all metazoan Orc1 orthologs possess an IDR and yet share little to no linear sequence similarity in this region. Despite their divergence, we demonstrate functional conservation of metazoan Orc1 IDRs from sponge to man. Collectively, these findings suggest that metazoan ORC implements a two-step chromatin binding mechanism, with IDR-dependent chromatin tethering necessarily precedeing and poising ORC for ATP-dependent DNA binding, which we propose occurs opportunistically as nucleosome free regions become available. More broadly, we provide evidence that IDRs with no sequence similarity can nonetheless be functionally conserved, and we develop the concept of compositional homology to explain this.

## RESULTS

### The Orc1 IDR is necessary and sufficient for regulated chromosome binding

We previously found that the ATP-binding and hydrolysis motifs of *Drosophila* Orc1 – motifs which are indispensable for ATP-dependent DNA binding *in vitro* (5,24) – are dispensable for *Drosophila* ORC’s chromatin recruitment *in vivo* (19). We therefore sought to identify the essential chromatin-recruitment element of *Drosophila* ORC. We have previously shown that Orc1 contains an Intrinsically Disordered Region (IDR) that in metazoans mediates DNA-dependent phase separation (19). This led us to ask whether the Orc1 IDR may also underlie chromatin recruitment. To test this, we generated transgenic fly lines expressing mNeonGreen (mNG) tagged full-length *Drosophila* Orc1 (mNG-Orc1), a construct lacking the IDR (mNG-Orc1^ΔIDR^), or the IDR alone (mNG-Orc1^IDR^). These transgenes were cloned from genomic DNA with endogenous promoter and stop sequences (**S1A-C Fig**) and transgene chromatin-binding dynamics were assessed by confocal fluorescence microscopy in live embryos 1.5-2 hrs after fertilization (nuclear cycles 10-13) (**Fig 1A-C**).

**Figure 1:**
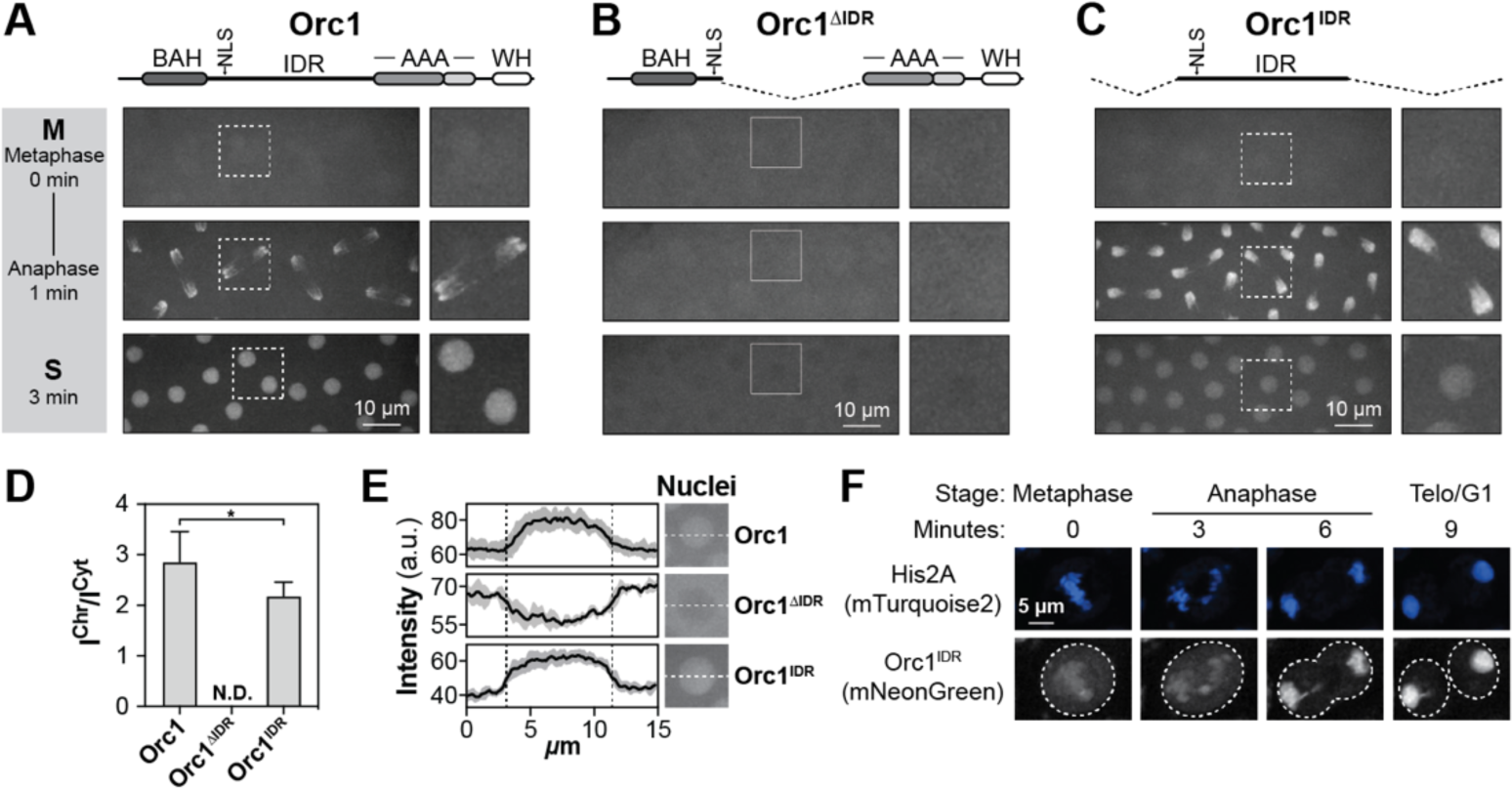
The fly Orc1 IDR is necessary and sufficient for chromatin recruitment in vivo. A) Fly embryos expressing an mNeonGreen-tagged full-length Orc1 transgene were imaged live and Orc1 chromatin binding dynamics assessed by confocal fluorescence microscopy during nuclear cycles 10-13. B) Chromatin binding dynamics of an mNeonGreen-tagged Orc1 deletion construct that lacks most of the IDR. C) Chromatin binding dynamics of an mNeonGreen-tagged Orc1 deletion construct that lacks all globular domains but retains the IDR. D) Ratio of mNeonGreen-Orc1 intensity on chromosomes versus the cytosol. No chromosome enrichement was observed for Orc1ΔIDR (N.D., not determined). A t-test was used to compare levels of chromosome enrichement (* = p < 0.05). E) Line intensity profile of mNeonGreen-Orc1 signal within and outside of the nucleus during S-phase. F) An mNeonGreen-tagged Orc1IDR was expressed in D. melanogaster S2 tissue cultures cells where we observed chromatin binding dynamics similar to embryos. Chromatin was visualized by co-expression of mTurquoise2-tagged His2A (blue).

We first assessed the dynamics of the full-length Orc1 transgene and found that mNG-Orc1 is recruited to chromatin in anaphase where it appears to uniformly coat chromosomes (**Fig 1A**). During anaphase, the chromosome intensity of mNG-Orc1 is 2.9-fold (± 0.60) above its cytosolic levels (**Fig 1D**). At this stage of *Drosophila* development, S-phase begins immediately after mitosis (i.e., there is no gap phase) and our results show that Orc1 remains enriched in the nucleus throughout this cell cycle phase (**Fig 1A**, bottom panel, and **Fig 1E**, top panel). Notably, the transition from S to M phase is evident from the loss of mNG-Orc1 signal in the nucleus that is the result of nuclear envelope breakdown upon mitotic entry (**Fig 1A**, compare bottom and top panels, and (19)). These results establish a baseline understanding of Orc1 cellular dynamics and are consistent with previous reports of ORC dynamics in the early embryo (19,25).

We next assessed the cellular dynamics of mNG-Orc1^ΔIDR^, a construct that lacks most of the predicted disordered region but retains the N-terminal nuclear localization signal (NLS). mNG-Orc1^ΔIDR^ was fully defective in recruitment to chromatin in anaphase (**Fig 1B**, middle panel) and, consequently, was depleted from the nucleus upon entry into S phase (**Fig 1B**, bottom panel, and **Fig 1E**, middle panel). Interestingly, in late S phase, mNG-Orc1^ΔIDR^ briefly transitions to a nuclear enriched state before it again disperses after nuclear envelope breakdown (**S1D Fig**). We are unsure how to interpret this change in nuclear enrichment except to say that mNG-Orc1^ΔIDR^ retains nuclear localization capabilities. Together, these data demonstrate that the Orc1 IDR is required for anaphase chromosome recruitment and provide a mechanistic rationale for our previous report that deletion of the Orc1 IDR is embryonic lethal (19).

We next assessed the *in vivo* chromatin binding dynamics of the Orc1 IDR alone (residues 187-549) to determine whether it is sufficient for chromatin recruitment. To test this, we prepared two transgenes: a genomic construct (mNG-gOrc1^IDR^) that spans part of two exons and the intervening intron, and a cDNA construct (mNG-Orc1^IDR^) (**S1C Fig**). No difference was observed in the chromatin binding dynamics of these two transgenes (compare **Fig 1C** and **S1E Fig**) and the data that follow derive from the cDNA construct. Despite lacking the globular domains which we generally associate with Orc1 function (e.g., BAH and AAA+ domains), the bulk cellular dynamics of mNG-Orc1^IDR^ were visually indistinguishable from that of the full-length protein (**Fig 1C**). Indeed, we find that mNG-Orc1^IDR^ is homogenously distributed as the embryo enters mitosis (**Fig 1C**, top panel), it then rapidly and uniformly binds chromosomes in anaphase (**Fig 1C**, middle panel), and remains nuclear enriched throughout S phase (**Fig 1C**, bottom panel, and **Fig 1E**, bottom panel). The anaphase chromosome intensity of mNG-Orc1^IDR^ is 2.2-fold (± 0.28) above its cytosolic level (**Fig 1D**), slightly lower than what we observed for the wild-type form of the protein. We also assessed Orc1^IDR^ dynamics in *D. melanogaster* tissue culture cells (S2 cells) to determine whether the function of this region is conserved in other developmental stages. We observed chromatin binding dynamics equivalent to what we observed in embryos, with Orc1^IDR^ showing loading onto anaphase chromosomes (**Fig 1F**). The only obvious distinction from embryos was a low level of Orc1^IDR^ binding to metaphase chromosomes. Together these data demonstrate that the Orc1 IDR is both necessary and sufficient for regulated association with mitotic chromosomes and provides a molecular explanation for ORC’s ATP-independent chromatin-binding capabilities (19).

### The Orc1 IDR mediates ATP-independent DNA binding in vitro

Recent studies show that ORC’s *in vitro* DNA binding activity requires ATP but not the Orc1 IDR (8,26). Paradoxically, we observe that ORC’s chromatin recruitment *in vivo* requires the Orc1 IDR but not ATP binding (**Fig 1** and (19)). To understand this discrepancy, we used fluorescence polarization to measure the *in vitro* DNA binding affinity of recombinant ORC complexes representative of the transgenes we produced for *in vivo* imaging (**Fig 1**), including the full-length ORC holocomplex (ORC), an ORC holocomplex lacking the Orc1 IDR (ORC^Δ1IDR^), and the isolated Orc1 IDR (Orc1^IDR^) (**S2A Fig**). ORC binds DNA in a sequence non-specific fashion (27) and we therefore used a random sixty basepair duplex DNA labeled with fluorescein (FITC-dsDNA) as a substrate in fluorescence polarization DNA binding assays.

We first assayed DNA binding under conditions similar to those used previously (8) and confirmed that ORC possesses high-affinity DNA-binding activity that is strictly dependent on ATP (**Fig 2A**, K_d_ = 7 nM ± 0.3). We reasoned that these assay conditions, which contain relatively high, non-physiological concentrations of potassium glutamate ([KGlut] = 300 mM), may selectively impede an IDR-dependent interaction with DNA to impose a dependency on ATP. We therefore repeated the DNA binding assay under the same conditions except using physiological levels of salt ([KGlut] = 150 mM KGlut) (**Fig 2B**). These conditions unveiled an ATP-independent mechanism of high-affinity DNA binding (**Fig 2B**, dotted line, K_d_ = 34 nM ± 10). The addition of ATP still modestly stimulated ORC’s affinity for naked DNA (**Fig 2B**, solid line, K_d_ = 6 nM ± 3).

**Figure 2:**
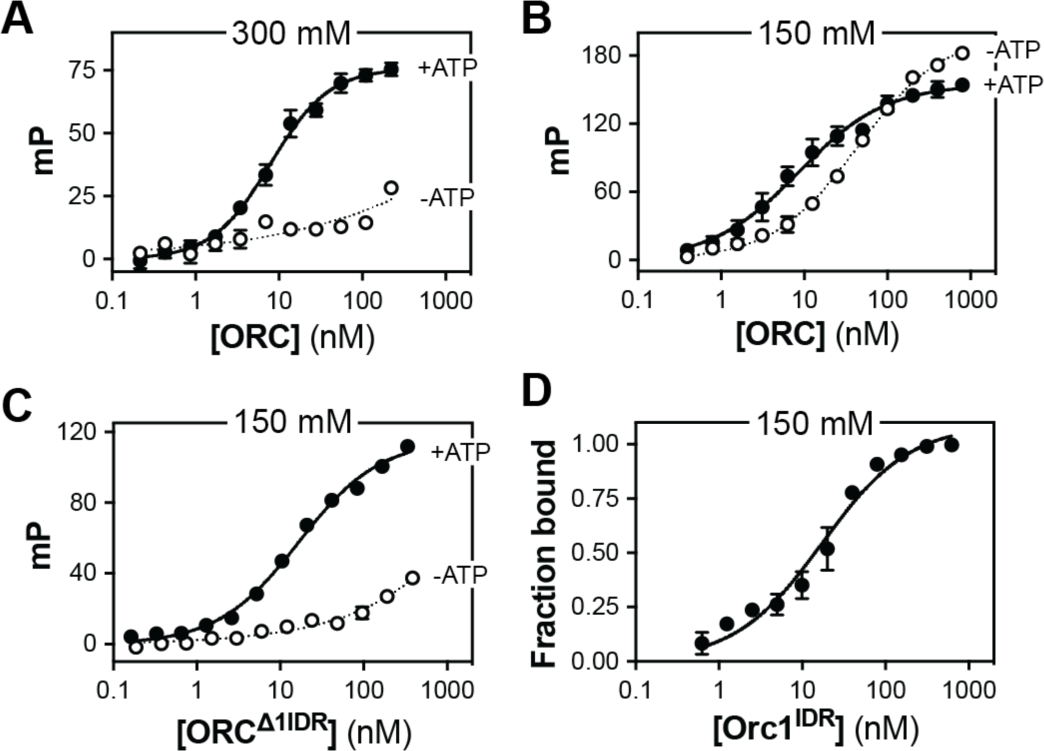
The Orc1 IDR facilitates ATP-independent DNA binding in vitro. A) A fluorescence polarization assay was used to measure the affinity of the ORC holocomplex for FITC-dsDNA in the presence (black line, K_d_ = 7 nM ± 0.3) and absence of ATP (dotted line, K_d_ = N.D.). Reaction conditions were 50 mM HEPES pH 7.5, 300 mM KGlutamate, 10% glycerol, 1 mM BME, 5 mM MgOAc (± 1 mM ATP). B) Same as A) except salt concentration was reduced to physiological (150 mM KGlutamate). ORC’s DNA binding affinity was assessed in the presence (black line, K_d_ = 6 nM ± 3) and absence of ATP (dotted line, K_d_ = 34 nM ± 10). C) The DNA binding affinity of an ORC holocomplex lacking the disordered region of Orc1 (ORCΔ1IDR) was assessed in the presence (black line, K_d_ = 19 nM ± 9) and absence of ATP (dotted line, K_d_ = N.D.) at 150 mM KGlutamate. D) Quantitation of an electrophoretic mobility shift assay (EMSA) used to measure binding of the isolated Orc1 IDR to Cy5-dsDNA in the absence of ATP (Orc1IDR, K_d_ = 15 nM ± 4).

To test whether the ATP-independent DNA binding we observe *in vitro* (**Fig 2B**) is mechanistically equivalent to the IDR-dependent chromatin recruitment we observe *in vivo* (**Fig 1**), we produced a mutant ORC holocomplex that lacks the Orc1 IDR (ORC^Δ1IDR^, **S2A Fig**) and assayed DNA binding. ORC^Δ1IDR^ retained high-affinity ATP-dependent DNA binding, with only a small reduction in affinity compared to the full-length complex (**Fig 2C**, solid line, K_d_ = 19 nM ± 9). Conversely, ATP-independent DNA binding by ORC^Δ1IDR^ was severely impaired (**Fig 2C**, dotted line, K_d_ > 1 µM). Finally, we produced the isolated Orc1 IDR (Orc1^IDR^, **S2A Fig**) and assessed its affinity for FITC-dsDNA by electrophoretic mobility shift assay (EMSA) (raw data shown in **S2B Fig**). We quantified the loss of free DNA with increasing concentrations of Orc1^IDR^ and found that it binds dsDNA with low nanomolar affinity (**Fig 2D**, solid line, K_d_ = 15 nM ± 4). These data demonstrate that ORC possesses two mechanistically separable DNA binding modes, one that requires the IDR and the other which requires ATP, and both are required *in vivo* (**Fig 1** and (19)).

### Phosphorylation of the Orc1 disordered region underlies regulated chromatin binding

Anaphase onset is coordinated with a cessation in Cyclin-dependent kinase (CDK) activity which, through an unknown mechanism, promotes ORC binding to chromosomes (25). This observation, together with our discovery of the DNA and chromatin binding activity of the Orc1 IDR, led us to hypothesize that phosphorylation of the Orc1 IDR may govern ORC’s chromatin binding dynamics. Consistently, the *D. melanogaster* Orc1 IDR possesses an abundance of CDK consensus motifs (‘[S/T]P’) (19,28). To test this idea, we generated stable *D. melanogaster* S2 tissue culture cell lines that express mTurquoise2-*Dm*Histone2A together with either mNeonGreen-tagged wild-type Orc1 (mNG-Orc1), the Orc1 IDR alone (mNG-Orc1^IDR^), or an Orc1 IDR variant where every CDK/Cyc consensus motif has been mutated ([S/T]P→AP, Orc1^IDR-ΔP^) (**Fig 3A**). We then used confocal fluorescence microscopy to assess the chromatin binding dynamics of each Orc1 transgene throughout mitosis. Each image set (blue and green channels) was thresholded for either mTurquoise2-*Dm*Histone2A (**Fig 3A**, blue) or mNeonGreen-Orc1 (**Fig 3A**, green) to create regions of interest (ROI) encompassing the chromosomes or cytosol, respectively, and the measured intensity of mNG within each ROI was used to calculate chromosome partitioning of Orc1 in metaphase versus telophase, the interval over which ORC loads onto chromosomes (**Fig 1**).

**Figure 3:**
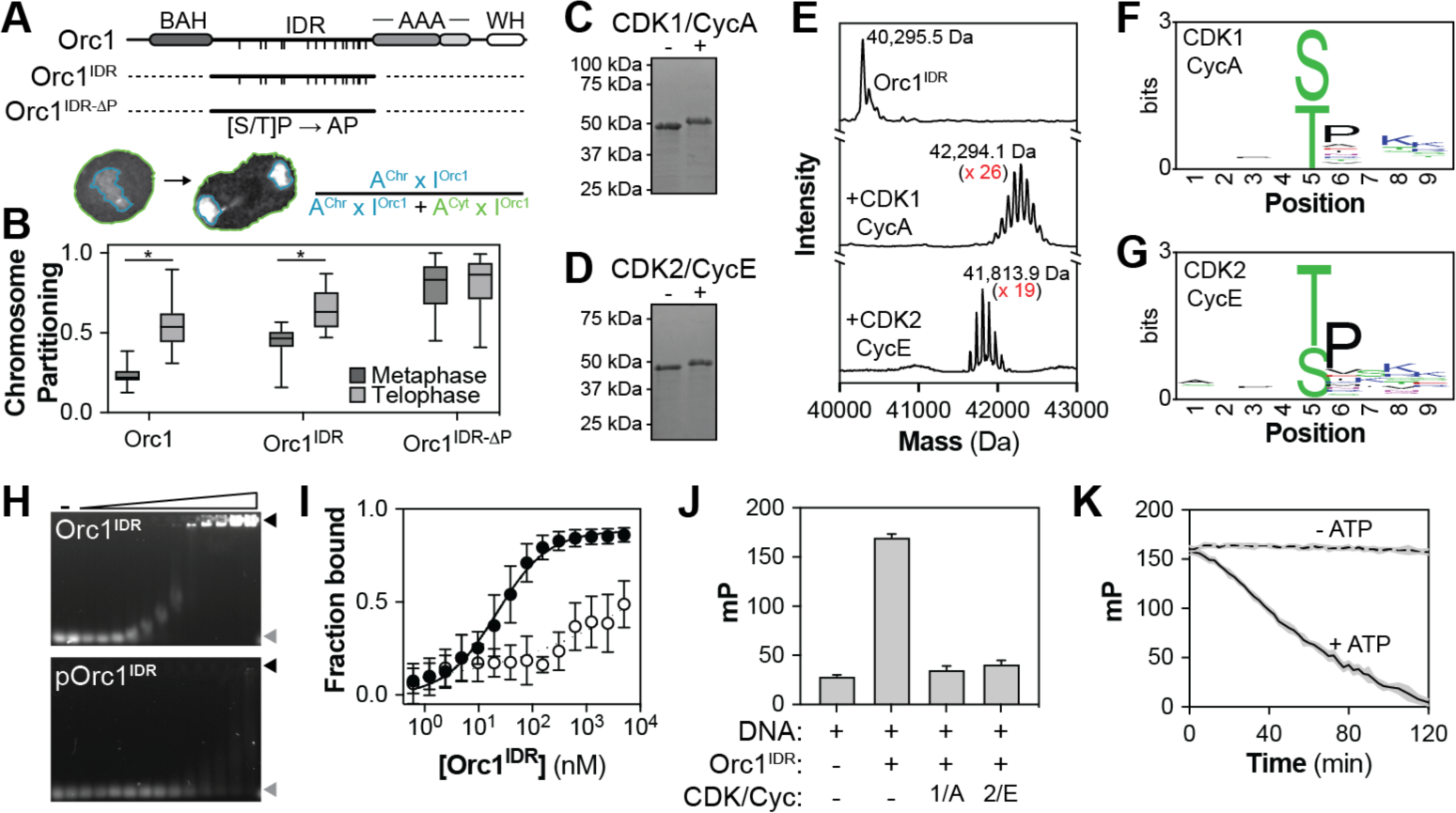
Phosphorylation of the Orc1 IDR regulates DNA and chromatin binding. A) Orc1 contains multiple putative CDK/Cyc phosphorylation sites within its IDR (indicated by hash marks). *D. melanogaster* S2 cell lines were generated that stably express full-length Orc1, the IDR alone, or an IDR variant that can no longer be phosphorylated. Each construct was tagged at the N-terminus with mNeonGreen, was co-expressed with mTurquoise2-His2A, and mitotic chromosome partitioning was measured in live cells. B) The chromosome partitioning of each Orc1 construct was assessed in metaphase and telophase. More than twenty cells were imaged for each construct and a t-test used to calculate significance (* = p < 0.05). C) *In vitro* reconstitution of Orc1^IDR^ phosphorylation (12 µM) by CDK1/CycA (2.4 µM) and D) CDK2/CycE (2.4 µM). E) Intact mass spectrometry was used to quantitate the level of Orc1^IDR^ (top, MW = 40,295.8 Da) phosphorylation induced by CDK1/CycA (middle) or CDK2/CycE (bottom). Indicated MW corresponds to highest intensity peak. F) Sequence logo of the CDK1/CycA-dependent phosphorylation sites identified by trypsin digestion of the phosphorylated samples and subsequent liquid chromatography mass spectrometry. G) Same as F), except for CDK2/CycE-phosphorylation of the Orc1^IDR^. H) The DNA binding affinity of unphosphorylated (Orc1^IDR^, top) and CDK2/CycE phosphorylated Orc1^IDR^ (pOrc1^IDR^, bottom) was measured by EMSA. I) Quantitation of H) (Orc1^IDR^ K_d_ = 21 nM and pOrc1^IDR^ K_d_ = N.D.). J) The purified Orc1^IDR^ (5 µM) was treated with either CDK1/CycA (1 µM) or CDK2/CycE (1 µM) in the presence of ATP (1 mM) and subsequently added to FITC-dsDNA (2 nM) and DNA-binding measured by fluorescence polarization. K) Time-resolved fluorescence polarization was used to assess how progressive phosphorylation of the Orc1^IDR^ impacts DNA binding. Orc1^IDR^ (5 µM) was premixed with CDK2/CycE (53 nM) and ATP (0 or 1 mM) added at time point = 0 at which point fluorescence polarization readings were begun.

Analysis of mNG-Orc1 chromosome partitioning revealed that, on average, 24% (± 6%) of the total Orc1 signal is associated with chromosomes in metaphase and that this more than doubles in telophase (56% ± 14%, **Fig 3B**). This is consistent with our *in vivo* results (**Fig 1**), except that Orc1 showed no binding to metaphase chromosomes in the early embryo. Similarly, mNG-Orc1^IDR^ showed a cell cycle-dependent increase in chromosome partitioning, increasing from 44% (± 10%) in metaphase to 65% (± 12%) in telophase (**Fig 3B**). While mNG-Orc1 and mNG-Orc1^IDR^ had overall similar dynamics, we note that the IDR alone showed significantly higher levels of chromosome partitioning in both metaphase and telophase, but why this occurs we do not currently understand. Finally, we assessed chromosome partitioning of mNG-Orc1^IDR-ΔP^, the Orc1 IDR variant that can no longer be phosphorylated. In striking contrast to mNG-Orc1 and mNG-Orc1^IDR^, mNG-Orc1^IDR-ΔP^ was highly enriched even on metaphase chromosomes (79% ± 15%) and showed no significant increase as the cells progressed into telophase (81% ± 15%, **Fig 3B**). These data demonstrate that in the absence of IDR phosphorylation Orc1 becomes constitutively associated with chromosomes.

We next sought to determine whether phosphorylation directly inhibits the IDR’s ability to interact with DNA. We therefore reconstituted IDR phosphorylation *in vitro* with the purified Orc1 IDR (Orc1^IDR^) in combination with CDK1/CycA or CDK2/CycE complexes purified from insect cells. In the presence of ATP, the addition of CDK1/CycA (**Fig 3C**) and CDK2/CycE (**Fig 3D**) resulted in a reduction of Orc1^IDR^ mobility on SDS-PAGE, strongly suggestive of phosphorylation. Similarly, treatment of the ORC holocomplex with kinase results in a noticeable shift of the Orc1 band as assessed by SDS-PAGE (**S3A Fig**). We confirmed IDR phosphorylation using intact mass spectrometry which also enabled quantitation of the number of sites phosphorylated by each kinase (**Fig 3E**). Although the Orc1 IDR has only fifteen putative CDK/Cyc consensus motifs, CDK1/CycA treatment added an average of 26 phosphate ions (**Fig 3E**, middle) and CDK2/CycE treatment added an average of 19 phosphate ions (**Fig 3E**, bottom). The precise phosphorylation sites were mapped by trypsin proteolysis followed by liquid-chromatography and mass spectrometry which revealed significant flexibility in CDK/Cyc motifs (**Fig 3F-G**). These data are visualized as a sequence logo derived from an alignment of the eight residues immediately surrounding each modified amino acid and, as expected, reveals a preference for a proline residue immediately C-terminal to the phospho-site (either Ser or Thr).

We next assessed the impact of phosphorylation on the Orc1 IDR’s *in vitro* DNA binding activity. We therefore repeated our purification of Orc1^IDR^ except integrated CDK2/CycE-dependent phosphorylation into our workflow and removed the kinase during the final size exclusion chromatography step. This resulted in a highly pure phosphorylated Orc1^IDR^ (pOrc1^IDR^) but with a slightly reduced level of phosphorylation (15 phosphates added, **S3B-C Fig**) compared to our previous assay executed at more concentrated reaction conditions (19 phosphates added, **Fig 3D-E**). We then used EMSAs to directly compare the DNA-binding affinity of Orc1^IDR^ and pOrc1^IDR^ (**Fig 3H-I**). Compared to the unphosphorylated sequence, pOrc1^IDR^ showed a >100-fold reduction in DNA binding affinity. This effect was confirmed for CDK1/CycA using fluorescence polarization DNA-binding assays (**Fig 3J**). Here, 5 µM Orc1^IDR^ was incubated with CDK1/CycA (1 µM) or CDK2/CycE (1 µM) in the presence of ATP before adding FITC-dsDNA and measuring fluorescence polarization. Notably, pre-treatment with either kinase markedly reduced the IDR’s DNA binding activity, suggesting that many CDK/Cyc pairs are likely sufficient to regulate Orc1.

Our experiments show that the Orc1 IDR is heavily phosphorylated by CDK/Cyc and raises the question of whether progressive phosphorylation events have a cooperative or linear impact on the IDR’s DNA binding activity. To address this, we premixed Orc1^IDR^ (5 µM), CDK2/CycE (50 nM), and FITC-dsDNA (2 nM) in the absence of ATP and placed the sample on ice. Subsequently, ATP was added, the sample was mixed, and fluorescence polarization readings were immediately and repeatedly taken for the phosphorylated sample (**Fig 3K**, solid line) and an unphosphorylated control (no ATP) (**Fig 3K**, dashed line). Over the course of the experiment (2 hours) we observed a linear reduction in fluorescence polarization in the presence of ATP but not in its absence. Assuming the rate of phosphorylation is linear, these data suggest that instead of acting as a binary switch, phosphorylation functions as a rheostat to tune the IDR’s affinity for DNA. These data may help explain why Orc1 partitions onto metaphase chromosomes in tissue culture cells but not in embryos (**Fig 1A,F**), two systems which may have a differential balance of kinase and phosphatase activity and therefore differing basal levels of IDR phosphorylation in metaphase. Altogether, these data suggest that regulatory CDK/Cyc-dependent phosphorylation blocks ORC activity by directly inhibiting the ability of the IDR to engage chromatin.

### The Orc1 IDR possesses multiple redundant DNA binding motifs

Our data show that the Orc1 IDR is an essential chromatin tethering element and we next sought to understand its mechanism of DNA binding. To identify functionally relevant regions, we assessed basic chemical features of the 363 amino acid long sequence, including the location of charged residues and the net charge per residue along the sequence (**Fig 4A**). This approach was motivated by our observation that ORC’s ATP-independent DNA binding activity, which is dependent on the IDR, is salt sensitive (**Fig 2A-C**). In general, the Orc1 IDR contains more basic residues than it does acidic (isoelectric point, pI = 10.1) and the charged residues do not appear to be organized in any obvious pattern. The C-terminal residues represent the most basic portion of the sequence (residues 487-549, pI = 11.5) and contains a cluster of basic residues (known as the basic patch, BP) that were previously shown to be important for ORC’s ATP-dependent DNA binding (8). We reasoned that the basic patch may likewise impart DNA binding activity to the IDR alone and we therefore purified an Orc1 IDR construct with a basic patch deletion (Δ520-533, Orc1^IDR-ΔBP^) and assayed its affinity for FITC-dsDNA by EMSA (**Fig 4B** and **S4A-B Fig**). Orc1^IDR-ΔBP^ bound dsDNA tightly (K_d_ = 8 nM ± 2) with no significant difference in affinity compared to the full-length IDR (**Fig 2D**, K_d_ = 15 nM ± 4).

**Figure 4:**
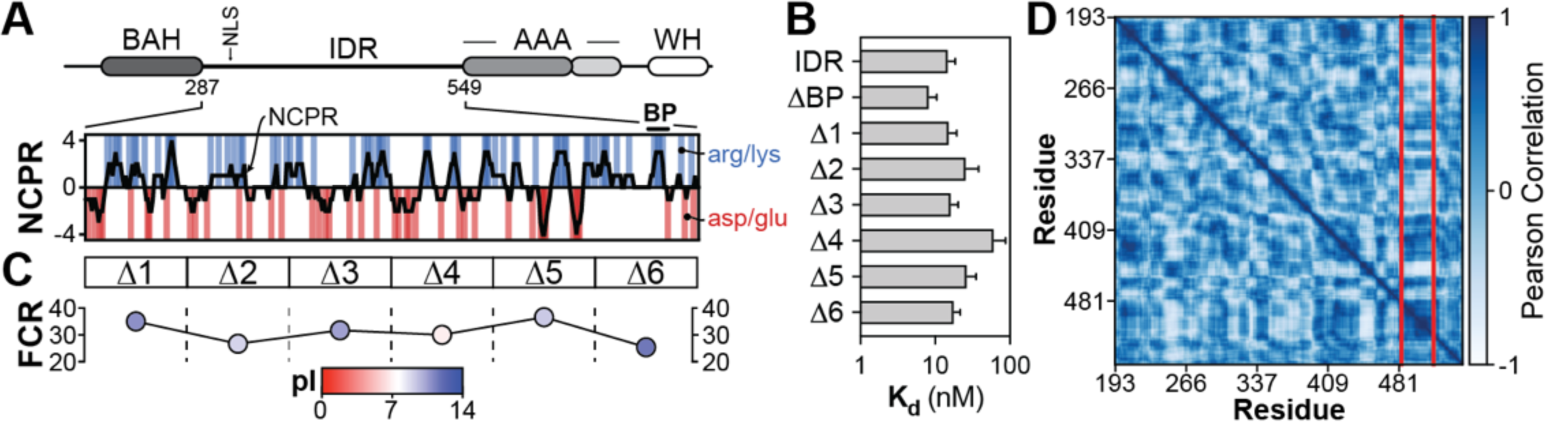
The Orc1 disordered region possesses multiple redundant DNA binding motifs. A) The domain architecture of DmOrc1 and a plot of the IDR’s net charge per residue (NCPR) and location of charged amino acids. “BP” denotes the previously identified basic patch in fly Orc1 (8). B) EMSAs were used to calculate the DNA binding affinity of Orc1 IDR variants. All variants bound DNA with a K_d_ < 100 nM. C) For each deleted region (Δ1-6), the fraction charged residues (FCR) was calculated and plotted. Each point is colored by the pI of the deleted region. D) Chi-score analysis (29) of Orc1 IDR modularity identifies a single compositionally-biased region.

Given the absence of other obvious sequence features targetable by mutagenesis, we took an unbiased, deletion mapping approach to identify DNA binding motifs within the IDR. We purified six IDR variants containing consecutive 60 amino acid deletions spanning the length of the IDR (**S4A Fig**, Orc1^IDR-Δ1^ – Orc1^IDR-Δ6^). Each deleted region has a relatively high fraction of charged residues (FCR, ranging from 0.25 to 0.37) and, except for the fourth region which is slightly acidic, all regions have a basic pI (ranging from 8.9 to 11.5) (**Fig 4C**). These analyses provided no obvious candidate DNA binding region(s) and when we measured the DNA binding affinity of each deletion construct, we found that all retained high-affinity DNA binding with a K_d_ < 100 nM (**Fig 4B** and **S4C-H Fig**). The only variant with a K_d_ significantly different from the full-length IDR was Orc1^IDR-Δ4^ which nonetheless still bound DNA with high affinity (K_d_ = 62 nM ± 26).

In a final effort to identify DNA binding regions, we analyzed the Orc1 IDR with a bioinformatic algorithm we have recently developed that uses the chi-square test statistic to quantify local variance in amino acid composition across a sequence. This approach led us to the unexpected discovery that many IDRs contain non-random, sequence-spanning compositional patterns that organize the sequence into juxtaposed modules of distinct compositional bias (29). However, unlike many other disordered regions (29), the fly Orc1 IDR is relatively unremarkable with weak modularity and no repetitive module types (**Fig 4D**). There are only two non-random boundaries (at a 95% confidence interval) that delimit a threonine rich region from a large N-terminal and shorter C-terminal module which both have high sequence complexity and no obvious sequence patterns. The threonine rich module is contained entirely in the region deleted in Orc1^IDR-Δ6^, which had no effect on DNA binding (**Fig 4B**). Altogether, these results afforded no further insight into the mechanism of DNA binding and suggest that the fly Orc1 IDR has a relatively uniform sequence landscape with highly redundant DNA binding sequence features scattered throughout its length.

### Metazoan Orc1 IDR orthologs have compositional homology but weak linear sequence similarity

We next assessed natural sequence variation amongst Orc1 IDR orthologs with the prediction that redundant DNA binding motifs would be conserved and that sequence alignments would divulge their location. We first compiled a list of twenty Orc1 orthologs selected from species representing many of the major metazoan phyla, including Porifera (sponges), Cnidaria (e.g., jelly fish), Arthropods (insects), Echinoderms (e.g., starfish), and Chordates, amongst others. The domain organization of each ortholog was predicted using Pfam (30) and Metapredict (31) to identify globular domains and regions of intrinsic disorder, respectively. Every Orc1 ortholog we assessed possessed both a AAA+ and winged helix (WH) domain (**Fig 5A**), regions which are known to be integral components of the ring-shaped ORC core complex conserved across eukaryotes (32). Each ortholog also possessed an IDR N-terminal to the AAA+ domain. In the sequences sampled, Orc1 IDRs were highly variable in length with the shortest being only 90 amino acids (Placozoa) and the longest nearly 500 amino acids (Echinoderm). Interestingly, both Echinoderm and Porifera Orc1 orthologs were found to possess AT-hook motifs embedded within their IDR. This DNA-binding motif interacts with AT-rich sequences and is the same motif that underlies chromatin binding of *S. pombe* ORC (14,15). Our alignments also revealed that Orc1’s Bromo Adjacent Homology (BAH) domain has been lost in certain phyla, including Tardigrades and Platyhelminthes (flatworms) (**Fig 5A**).

**Figure 5:**
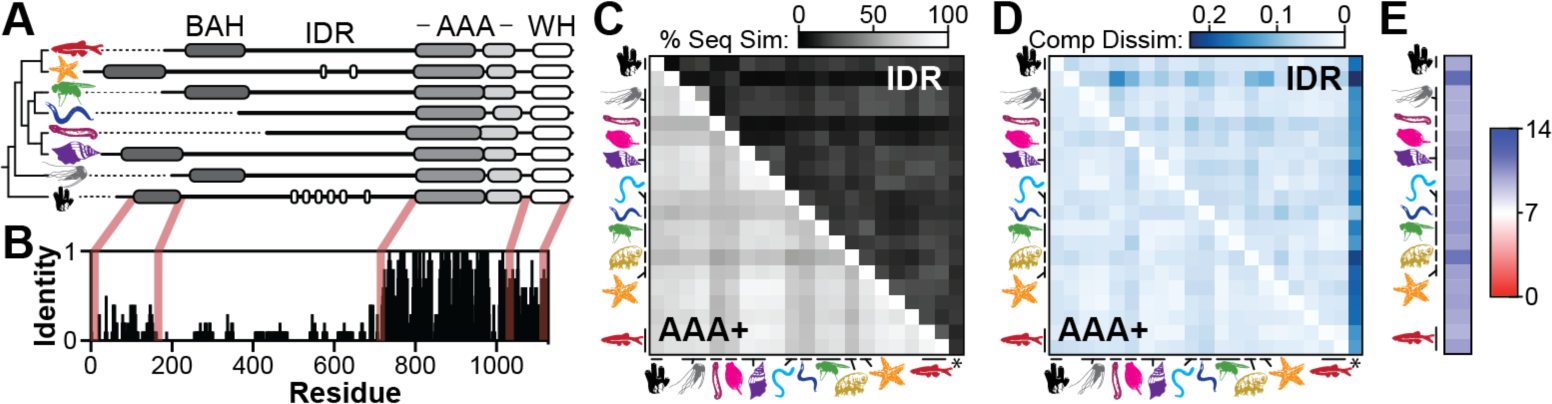
Metazoan Orc1 IDR orthologs have compositional homology but lack linear sequence similarity. A) The domain architecture of Orc1 is largely conserved across metazoans. Represented phyla include (from top to bottom): Chordate, Echinoderm, Arthropod, Nematode, Platyhelminthe, Mollusc, Cnidaria, and Porfiera. Echinoderm and Porifera Orc1 orthologs possess AT-hook motifs (white regions) embedded within their IDR. B) Sequence identity across twenty Orc1 orthologs. C) Pairwise comparison of linear sequence similarity between twenty Orc1 AAA+ and IDR orthologs. “*” column indicates a comparison between each Orc1 IDR and a randomly scrambled version of itself. D) Pairwise comparison of compositional dissimilarity between twenty Orc1 AAA+ and IDR orthologs. Lighter colors indicate sequences with similar sequence composition. “*” column indicates a comparison between each Orc1 IDR and a “normal” sequence (5% of each amino acid). E) All metazoan Orc1 IDRs assessed have a basic pI.

Using multiple sequence alignment, we measured the sequence identity across the Orc1 gene (**Fig 5B**). This analysis showed that the AAA+ domain is the most highly conserved region within Orc1, followed by the WH domain and then the BAH domain. Interestingly, our alignments failed to identify regions of sequence similarity within the disordered region. While it is known that IDRs evolve more rapidly than globular sequences (33), we were expecting some level of conservation based on the essential role this region plays in flies. We reasoned that the vast evolutionary time scale represented in our analysis (approximately 800 million years (34)) may mask conserved regions and we thus assessed pairwise sequence similarity for both the AAA+ domain and IDR (**Fig 5C**). The AAA+ domain (bottom left of diagonal) was highly conserved for each pair of sequences, possessing 54-94% sequence similarity. Conversely, the IDR (top right of diagonal) showed such weak conservation (4-55% sequence similarity) that in many cases it was unclear if the alignments revealed significant levels of homology or simply spurious registration of short sequences. To answer this, we generated alignments of each Orc1 IDR ortholog with a randomly scrambled version of itself (column marked with an asterisk) and found that sequence similarity ranged from 10-19%. Of the 190 pairs of Orc1 IDR orthologs, 82 have a score at or below that observed for the randomly scrambled sequences suggesting that in many cases the Orc1 IDR is so highly diverged that no sequence similarity remains.

The sequence divergence amongst Orc1 IDRs failed to advance our search for DNA binding motifs and further suggested that the function of the Orc1 IDR may not be conserved. Another intriguing possibility, however, is that the function of the Orc1 IDR does not depend on the linear ordering of amino acids, and instead relies only on sequence composition. We therefore assessed whether Orc1 IDRs are *compositionally homologous* using a newly developed metric that quantitatively compares the fractional composition of amino acids between sequences (29). The results were scaled such that sequences with the same composition have a score of 0 and sequences that have no shared amino acids have a score of 1. We calculated compositional homology between orthologous Orc1 AAA+ domains (**Fig 5D**, bottom left) and orthologous Orc1 IDRs (**Fig 5D**, top right). As a comparison, we calculated compositional homology between each IDR and a standardized sequence composed of 5% of each of the twenty amino acids (column marked with an asterisk). From this analysis it was clear that Orc1 IDR orthologs have a sequence composition much more similar to one another than to the standardized sequence, and that their level of compositional homology is, in many cases, on par with what is observed between AAA+ domains. These data suggest that the function of Orc1 IDRs could be conserved despite an absence of traditional forms of sequence homology.

### The function of the Orc1 IDR is conserved across metazoans

The key functional features of the *D. melanogaster* Orc1 IDR are its ability to bind DNA and chromatin (**Fig 1-3**) and we therefore set out to test whether Orc1 IDR orthologs share these functionalities. We synthesized the Orc1 IDR coding region from multiple metazoan organisms representing a diverse set of phyla and we were able to successfully express and purify the Orc1 IDRs from *A. millepora* (Cnidaria), *B. plicatilis* (Rotifera), *M. yessoensis* (Mollusca), *D. gyrociliatus* (Annelida), *H. dujardini* (Tardigrada), and *A. queenslandica* (Porifera). These IDRs range in size from 158-454 amino acids and all purified as a single band on SDS-PAGE except for *A. queenslandica* which was a mixture of full-length and proteolyzed product (**S6 Fig**). To assay DNA binding, we combined each IDR (600 nM) with FITC-dsDNA (2 nM) and assessed binding by EMSA (**Fig 6A**). For comparison we also assayed DNA-binding with the fly Orc1 IDR (*Dm*Orc1^IDR^) and the mutated variant that lacks phosphorylation sites (*Dm*Orc1^IDR-ΔP^). At this concentration, each of the Orc1 IDRs caused a near complete loss of free FITC-dsDNA and we observed a heterogenous mixture of either well-shifted or smeared IDR•DNA complexes, indicative of variable stoichiometry or possibly a dynamic complex. Despite possessing AT-hook motifs, which are *bona fide* DNA binding elements, *Aq*Orc1^IDR^ appeared to have the weakest DNA binding. We suspect this is due to incompatibility of our probe DNA (which has 50% AT content) and the specificity of AT-hook motifs for AT-rich regions of DNA.

**Figure 6:**
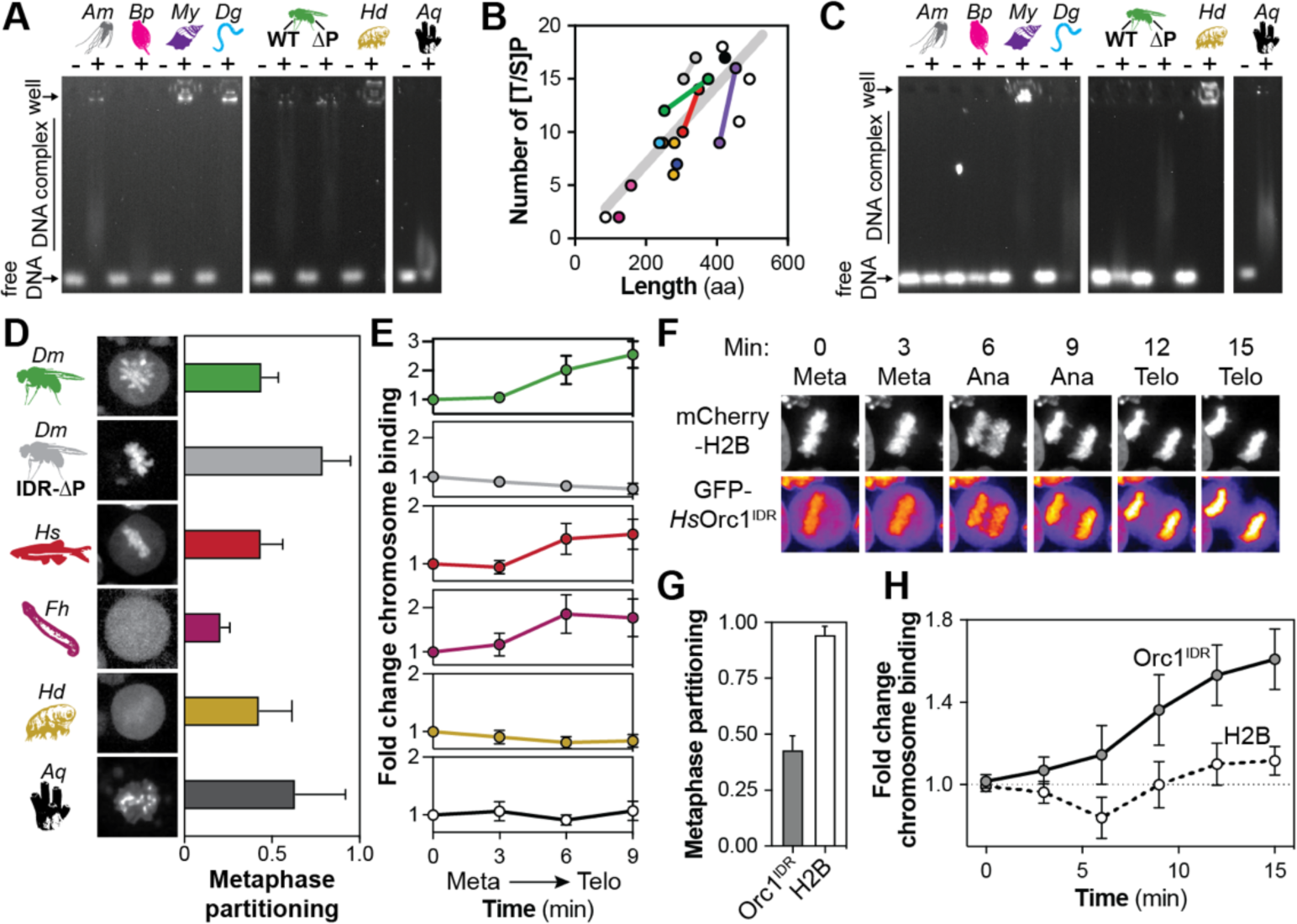
The Orc1 IDR is functionally conserved across metazoans. A) EMSAs were used to assess binding of Orc1 IDR orthologs (600 nM) to Cy5-dsDNA (2 nM). Orthologs from the following phyla were used: Cnidaria (Am), Rotifer (Bp), Mollusc (My), Annelid (Dg), Arthropod (Dm), Rardigrade (Hd), and Porifera (Aq). ‘ΔP’ is the DmOrc1 IDR with mutated phosphorylation sites (all “[S/T]P” (“AP”). B) Graph showing the number of phosphorylation sites (“[T/S]P”) versus the length of twenty Orc1 IDR orthologs. Each point represents an IDR and is colored by phyla. IDRs from the same phyla are connected by a line. The grey line indicates the line of best fit. C) Each Orc1 IDR ortholog (600 nM) was phosphorylated by CDK2/CycE and DNA binding assessed by EMSA. D) Metaphase images of Drosophila S2 cells expressing Orc1 IDR orthologs tagged with mNeonGreen. Orthologs from the following phyla were used: Arthropod (Dm), Chordate (Hs, human), Platyhelminthe (Fh), Tardigrade (Hd), and Porifera (Aq). E) Live imaging experiments in Drosophila S2 cells were used to calculate the fold change in mitotic chromosome recruitment for each Orc1 IDR ortholog. F) HeLa cells (stably expressing mCherry-H2B) were transiently transfected with GFP-tagged human Orc1IDR (HsOrc1IDR) and live imaging used to assess HsOrc1IDR recruitment to mitotic chromosomes. G) Metaphase partitioning of HsOrc1IDR and H2B. H) Live imaging experiments were used to calculate the fold change in chromosome recruitment of HsOrc1IDR and H2B throughout HeLa cell mitosis.

We next asked whether the DNA binding activity of metazoan Orc1 IDRs is regulated by phosphorylation. First, we assessed if CDK/Cyc sites (“[S/T]P”) are present in metazoan Orc1 orthologs and found that the number of sites is linearly correlated with the length of the IDR (**Fig 6B**). Next, we combined purified *D. melanogaster* CDK2/CycE (0.5 µM) with each IDR (2.5 µM) and, after a 30-minute incubation, we incubated each IDR (600 nM) with FITC-dsDNA (2 nM) and assessed binding by EMSA (**Fig 6C**). As expected, treatment with CDK2/CycE inhibited the DNA binding activity of the *Dm*Orc1^IDR^ – as evidenced by the presence of a free DNA band – but not the variant with mutated phosphorylation sites (*Dm*Orc1^IDR-ΔP^). Interestingly, we found that about half of the purified Orc1 IDR orthologs are regulated by CDK2/CycE-dependent phosphorylation (including *Am*, *Bp*, and *Dm*), and the other half are clearly not regulated (including *My, Hd*, and *Aq*) or are only partially (*Dg*). One possible explanation for the observed differences is that the *Drosophila* CDK2/CycE complex may be effective against only a subset of our sequences. Alternatively, some sequences may legitimately lack regulation, such as *Hd*Orc1^IDR^, which has multiple minimal CDK/Cyc recognition motifs (“[S/T]P”) but only a single optimal site (“[S/T]PX[R/K]”). Likewise, we suspect that the more sequence specific DNA binding facilitated by the AT-hook motifs of *Aq*Orc1^IDR^ may be insensitive to phosphorylation. Collectively, these data indicate that despite an absence of sequence similarity, metazoan Orc1 IDRs universally possess the ability to bind DNA and suggest that phospho-regulation is broadly but not universally conserved.

The conservation of DNA binding activity prompted us to assess whether Orc1 IDR orthologs can also facilitate chromatin tethering within the cell. Testing this in each organism individually would be impractical and we therefore assessed functionality in a heterologous system. Specifically, we generated stable *D. melanogaster* cell lines (S2 cells) that co-express mNeonGreen-tagged Orc1 IDR orthologs alongside mTurquoise2-tagged Histone2A (for chromosome visualization). We were successful in generating stable cell lines for some Orc1 IDRs but not others, resulting in data sets on six orthologs (*Hs*Orc1^IDR^ (Chordate), *Fh*Orc1^IDR^ (Plateyhelminthes), *Hd*DmOrc1^IDR^ (Tardigrade), *Aq*Orc1^IDR^ (Porifera), *Dm*Orc1^IDR^ (Arthropod), and *Dm*Orc1^IDR-ΔP^), four of which are represented in our *in vitro* DNA binding assays. Each cell line was individually imaged by spinning disk confocal fluorescence microscopy and we collected still images to assess chromosome partitioning in metaphase (**Fig 6D**) as well as time series to examine if orthologs show regulated recruitment to chromatin as cells progress through mitosis (**Fig 6E**). The *Dm*Orc1^IDR^ (regulated chromatin binding) and *Dm*Orc1^IDR-ΔP^ (constitutive chromatin binding) data are derived from **Fig 3A-B** and are presented for comparison with other orthologs. As observed for the fly Orc1 IDR (metaphase partitioning = 44%), *Hs*Orc1^IDR^ (Chordate) was found to be moderately enriched on metaphase chromosomes (**Fig 6D**, red, metaphase partitioning = 44%) and showed enhanced chromatin binding as cells progressed through mitosis (**Fig 6E**, red). *Fh*Orc1^IDR^ (Platyhelminthes) did not appear visibly enriched on nor excluded from metaphase chromosomes (**Fig 6D**, purple, partitioning = 21%) but progression into the later stages of mitosis resulted in significant chromosome recruitment (**Fig 6E**, purple). There was relatively high cell-to-cell variability in the chromatin binding dynamics of *Hd*Orc1^IDR^ (Tardigrade, **Fig 6D**, gold, partitioning = 43%) and *Aq*Orc1^IDR^ (porifera, **Fig 6D**, black, partitioning = 63%) but on average they were moderately enriched on chromatin in metaphase and in both cases we observed no change in chromosome partitioning as the cells progressed through mitosis (**Fig 6E**, gold and black). Notably, *Aq*Orc1^IDR^ possesses a puncate distribution on chromosomes which we speculate is pericentromeric heterochromatin on the basis that the Chromosomal protein D1, the factor which normally occupies these chromosomal loci, also has multiple AT-hook motifs. The lack of regulation we observed for *Hd*Orc1^IDR^ and *Aq*Orc1^IDR^ is consistent with our *in vitro* data where we observed DNA binding regardless of CDK2/CycE phopshorylation (**Fig 6A-C**). Consistent with our *in vitro* DNA binding studies, these data suggest that chromatin binding is a universally conserved function of metazoan Orc1 IDRs, and that some othologous seqeunces likely mediate a constitutive association with chromatin (e.g., *Hd* and *Aq* Orc1 IDRs).

As a final test of the functional conservation of Orc1 IDRs, we assayed the chromatin binding dynamics of the human Orc1 IDR (*Hs*Orc1^IDR^) in human tissue culture cells. We constructed a transgene that expresses the human Orc1 IDR coding region fused at its C-terminus to GFP and transiently transfected the construct into HeLa cells stably expressing mCherry-tagged histone H2B (35). Cells were plated in glass bottom dishes which were scanned by confocal fluorescence microscopy for mitotic cells. The position of each mitotic cell was marked and subsequently imaged with an automated imaging routine that collected a z-stack every three minutes through mitosis (**Fig 6F**, representative image set). As seen in *D. melanogaster* S2 cells, *Hs*Orc1^IDR^ was moderately enriched on metaphase chromosomes (**Fig 6F-G**, metaphase partitioning = 44%) and, as cells progressed through mitosis, chromosome partitioning of the human Orc1 IDR increased to 1.6-times the level seen in metaphase (**Fig 6H**). We also quantitated mCherry-H2B dynamics through mitosis and, as expected, protein intensity remained essentially unchanged, being fully partitioned onto chromatin regardless of mitotic stage (**Fig 6F-H**). These data demonstrate that the Orc1 IDR is a DNA and chromatin binding element and that it is functionally conserved across the metazoan lineage.

## DISCUSSION

We report here that the fly Orc1 IDR is necessary and sufficient for *in vivo* chromosome recruitment of ORC and that CDK/Cyc-dependent multisite phosphorylation inhibits chromatin binding. These findings provide the molecular logic behind metazoan ORC’s regulated association with chromatin. Together with our previous work (19), these data suggest that chromatin is recalcitrant to ORC’s ATP-dependent DNA binding and encirclement activity, and we suggest that this necessitates chromatin tethering by the Orc1 IDR. Interestingly, we find that metazoan Orc1 IDR orthologs share little to no sequence similarity but nonetheless have a conserved functionality. We propose that a similar amino acid composition is sufficient to maintain the function of Orc1 IDR orthologs. These findings revise our understanding of the mechanism of metazoan DNA replication licensing and, more broadly, expand the relationship between a protein’s primary structure and function with the concept of compositional homology.

Over thirty years ago *S. cerevisiae* ORC was identified on the basis of its ATP-dependent association with yeast origins of replication (36). Subsequent studies found that human and fly ORC also possess ATP-dependent DNA binding, with the caveat that non-specific, nucleotide-independent interactions with DNA do occur (5,7,20). These data led to the notion that metazoan ORC, like *Sc*ORC, is recruited to chromatin via ATP-dependent DNA binding. Recent studies support this, showing that ATP is strictly required for fly ORC’s *in vitro* DNA binding activity (8,26). We have now shown that ATP binding is dispensable for chromatin recruitment of ORC *in vivo* (19) and that the metazoan Orc1 IDR is the essential chromatin tethering element (**Fig 1**). These findings present an apparent paradox with the reported ATP-dependence of ORC’s *in vitro* DNA binding activity (8). Our study rationalizes this discrepancy with *in vitro* experiments showing that ATP is only required for ORC’s DNA binding activity when salt concentrations are relatively high ([KGlutamate] = 300 mM). Indeed, at physiological salt concentrations (150 mM) we observe high-affinity, *ATP-independent* DNA binding by ORC, and this requires the Orc1 IDR (**Fig 2**).

Our results suggest that the mechanism of metazoan ORC chromatin binding is more akin to *S. pombe* than it is to *S. cerevisiae*. Indeed, the metazoan Orc1 IDR seems to be the functional analog of the AT-hook-containing *Sp*Orc4 N-terminus in the limited sense that both are required for chromatin tethering (14,15). This raises the question of why chromatin tethering is needed at all, since ORC’s ATP-dependent DNA binding and encirclement activity is clearly conserved across eukaryotes and is sufficient for high-affinity DNA binding *in vitro* (**Fig 2** and (26)). We propose that in metazoans chromatin is inherently restrictive to ATP-dependent DNA binding, and that this necessitates a chromatin tethering mechanism. This idea is most clearly supported by the observation that ORC^Δ1IDR^ still possesses high-affinity ATP-dependent binding to naked DNA *in vitro* (**Fig 2**) and yet deletion of the Orc1 IDR *in vivo* is lethal (19) and the protein can no longer bind chromatin (**Fig 1**). In hindsight, this conclusion appears self-evident, for we know that ORC, which requires at least forty base pairs of naked DNA for ATP-dependent binding (8), is ineffective at displacing nucleosomes (37,38). Further, internucleosomal linker DNA is relatively short (25-50 base pairs on average (39,40)) and occluded by histone H1 (41). Thus, the lack of naked DNA *in vivo* precludes ATP-dependent DNA binding and encirclement by ORC. This problem is further emphasized when one considers the nearly one hundred base pairs needed for full Pre-RC assembly (42). It seems that this problem is evaded altogether by *Sc*ORC as it is both constitutively associated with chromatin and binds specific DNA sequences that are maintained in a nucleosome free state (43). For these reasons we think an accessory chromatin tethering appendage is required for *S. pombe* and metazoan ORC (and likely most other eukaryotes) but not the *S. cerevisiae* complex.

Of course, ATP-dependent DNA binding and encirclement must and does occur within the context of the chromosome, and we think that chromatin tethering promotes this. We propose that ORC is first non-specifically and dynamically tethered to chromatin via the Orc1 IDR and that this state, though not a structurally resolvable Pre-RC intermediate, positions ORC to opportunistically bind and encircle nucleosome free regions in an ATP-dependent fashion. Thus, metazoan ORC does not select origins per se, but rather exploits the chromatin remodeling capabilities of other chromatin contextualized processes with which it has no direct connection. In line with this, genomics studies demonstrate that ORC is enriched at transcription start sites and other regions of high nucleosome turnover (44–46). This mechanism would endow the licensing machinery with an inherent flexibility to adapt to the cell-type specific transcriptional programs of multicellular organisms. We suspect that the DNA-binding IDRs of metazoan Cdt1 and Cdc6 (19) play a similar role by maintaining relatively high concentrations of these factors on chromatin to enable the efficient assembly and recycling of Pre-RC components, and this may be further supported by the ability of these proteins to phase separate with DNA (19,47).

While this work clearly demonstrates that the Orc1 IDR tethers ORC to chromatin, precisely how it does this remains to be determined. Our results show that the Orc1 IDR’s DNA binding motifs are highly redundant (**Fig 4**) and that Orc1 IDR orthologs retain DNA and chromatin binding capabilities (**Fig 6**) but, oddly, have little to no sequence similarity (**Fig 5**). Orc1 IDR orthologs do, however, possess a similar amino acid composition and all have a basic pI (**Fig 5**). We propose the term compositional homology to describe such sequence sets and our newly developed chi-score metric represents a quantitative means of classifying these (29). Simply put, these results suggest that the precise ordering of amino acids in the Orc1 IDR is not important and that these sequences likely interact non-specifically with the DNA backbone. Consistently, the Cdt1 IDR, which is compositionally homologous to the Orc1 IDR (19), retains the ability to bind DNA even when randomly scrambled (48). The sponge Orc1 IDR is somewhat unique in that it contains seven AT-hook motifs (e.g., *Aq*Orc1^473-482^: ‘RKR<u>GRP</u>RKEE’), the very same DNA-binding element found in the *Sp*Orc4 N-terminus (15,16). Interestingly, degenerate AT-hook motifs are also present in echinoderm (501-KQ<u>GRP</u>KK-507 and 564-RKR<u>GRP</u>RSVKK) and fission yeast (226-RGR<u>GRP</u>RK-233) Orc1 IDRs, suggesting the intriguing possibility that an AT-hook containing IDR was the ancient ancestral sequence from which Orc1 IDRs have diverged and have maintained DNA binding capabilities but with reduced specificity.

The Orc1 IDR’s non-specific, electrostatic-based interactions with DNA raise the question of how specific binding to DNA/chromatin is achieved. We have shown in a previous study that metazoan licensing factors undergo DNA-dependent phase separation, that their IDRs are required for this (19), and, at least for Cdt1, that other polyanions (RNA and poly-glutamate) function equally well as duplex DNA in stimulating phase separation (48). We therefore cannot rule out the possibility that, in addition to DNA, the Orc1 IDR is targeted to chromatin through interactions with chromatin-associated RNAs (49) and proteins containing an acidic surface (e.g., the H2A/H2B acidic patch), or a combination of these. In fact, we favor the idea that the IDR can interact with many different chromatin features which may provide context-independent chromatin association to support genome-wide licensing. In support of this, it has been shown that Orc1, and specifically the Orc1 IDR, interacts with chromatin-localized RNA and that this is important for both normal cellular replication as well as for licensing of viral genomes (50–52). How then is the IDR targeted specifically to chromatin and not to other RNA-enriched structures? This question remains to be answered but one interesting possibility is that IDR dephosphorylation is spatially restricted to the chromosome surface through chromatin-associated phosphatases, such as the RepoMan-Protein Phosphatase 1 (PP1) complex which, like ORC, binds chromatin at anaphase onset (53).

This study, in addition to our previous work with Cdt1 (48), adds the metazoan licensing factor IDRs to a short list of intrinsically disordered sequences whose function is known to rely solely on sequence composition and not on the linear ordering of amino acids. Other proteins in this category include linker histone H1 which retains the ability to compact DNA even when the sequence is randomly scrambled (54) and the prion domains of yeast Sup35p and Ure2p which can be scrambled without losing the ability to induce amyloid formation (55,56). Collectively, this limited set of experiments suggest that the function of certain IDR classes depends only on the combined attributes of length and fractional content of amino acids, and that functionally homologous IDRs could be identified by compositional homology alone. An interesting future direction is to extend these observations to other sets of IDR orthologs to compare the extent of compositional homology versus linear sequence similarity, which would likely provide important mechanistic insight into how function is encoded in disordered sequences.

In conclusion, this work demonstrates that metazoan ORC engages chromatin in a two-step process, with IDR-dependent chromatin tethering preceding ATP-dependent DNA binding and encirclement. We propose that chromatin tethering is likely a general solution to overcome the restrictive nature of chromatin, and evolution appears to have implemented this in various ways, including with the metazoan Orc1 IDR, the AT-hook motifs of *S. pombe* Orc4, and possibly hitherto unidentified mechanisms, such as the predicted Zn-finger in *Arabidopsis* Orc1. Our data suggest that the function of metazoan Orc1 IDRs does not depend on the linear ordering of amino acids and we provide evidence that amino acid composition alone is their defining feature. We anticipate that compositional homology will be an important concept for understanding the functional conservation of other classes of disordered domains that show limited or no linear sequence similarity.

## MATERIALS AND METHODS

### Generation of transgenic fly lines

The full-length *D. melanogaster* Orc1 (*Dm*Orc1) gene was cloned from genomic DNA (OregonR fly line) by PCR amplification of the protein coding regions plus 1 kilobase pair of regulatory sequence up and downstream of the start and stop codons. This sequence was inserted into vector pattB and Gibson Assembly was used to replace the start codon with the coding sequence for the fluorescent protein mNeonGreen. This construct was used as a template for around-the-horn mutagenesis to generate an IDR deletion construct (Δ249-541, DmOrc1^ΔIDR^). We generated two transgenes that express the IDR alone, one that was PCR amplified from genomic DNA and spanned two exons (Exon 1 and Exon 2) and one that was PCR amplified from cDNA. These constructs encode residues 187-549 of the Orc1 gene and were inserted into pattB with an N-terminal mNeonGreen tag. These constructs (pattB-mNG-g*Dm*Orc1, pattB-mNG-g*Dm*Orc1^ΔIDR^, pattB-mNG-g*Dm*Orc1^IDR^, pattB-mNG-c*Dm*Orc1^IDR^) were then injected into embryos for site specific PhiC31 integration into chromosome 3 at position attP2:68A4 (Genetivision Corporation). The resulting transgene lines were balanced and crossed to produce homozygous lines. Genomic DNA from homozygous lines was used as a template for PCR amplification of the inserted transgenes which were then confirmed by sequencing.

### Fluorescent imaging of *D. melanogaster* embryos

Homozygous transgenic fly lines were amplified and combined in bottles (200 – 500 flies per bottle) that were inverted onto agar plates smeared with yeast paste to induce egg laying. The collected embryos were dechorionated in 100% bleach for 2 min followed by extended washing with H_2_O. The embryos were then transferred into µ-slide 4-well glass bottom imaging dishes (Ibidi), covered with halocarbon oil, and then imaged. Images were acquired on a Nikon Ti2E microscope equipped with a Yokogawa CSU X1 spinning disc using a 60x oil immersion objective with the appropriate filter set and images collected with 488 nm laser power set to 12.7% and a 200 ms exposure. Z-stacks were collected at 1 µm intervals through the full embryo volume. Z-stacks were acquired every 30 seconds until nuclear cycle 14. Image processing was done with FIJI. For each image set, a maximum intensity projection was generated and the measured mNeonGreen-Orc1 intensity was used to calculate the ratio between Orc1 signal on chromosomes versus the cytosol. These quantitations required knowing when a cell was in a mitosis, which was evident from the doubling of nuclei.

### Generation of transgenic *D. melanogaster* S2 cells

To assess Orc1 cellular dynamics in *Drosophila* S2 cells, plasmids were constructed by cloning Orc1 and Orc1 IDRs into our custom pCopiaFP(mNG) vector which appends inserts with an N-terminal mNeonGreen tag. Coding regions included: full length *Drosophila* Orc1, *Drosophila* Orc1^IDR^ (residue 187-549), *Drosophila* Orc1^IDR-P-dead^ (residue 187-549 with every ‘[S/T]P’ mutated to ‘AP’) and metazoan Orc1^IDR^ orthologs (human Orc1 residues 177-484, *F. hepatica* (Platyhelminthes) Orc1 residues 1-267, *H. dujardini* (Tardigrade) Orc1 residues 1-278, and *A. queenslandica* (Porifera) Orc1 residues 177-593). All cloning was carried out using Ligation Independent Cloning (LIC). To visualize chromatin, *Dm*Histone2A was fluorescently labeled at its N-terminus with mTurquoise2 by cloning into our custome pCopiaFP(mTurquoise2) vector Subsequently, individual Orc1 constructs were co-transfected with vectors p8HCO (providing a methotrexate resistant gene) and pCopiaFP(mTurquoise2)-*Dm*His2A into *Drosophila* S2 cells (Expression Systems) maintained at 27°C in ESF 921 medium (Expression Systems). For transfection, S2 cells were seeded in 6-well plates at a density of 2 × 10^6^ cells/well. After 24 hr the medium was removed and supplemented with fresh medium. Transfections were carried out using Effectene reagent following the manufacturer’s protocol (Qiagen). Fourtyeight hours post transfection, the medium was replaced with fresh medium supplemented with 0.1 µg/ml methotrexate. Subsequently, the stably transfected cells were selected by replacing medium with methotrexate supplemented fresh medium every 3 days for 5 weeks.

### Fluorescent imaging of *D. melanogaster* S2 cells

Stably transfected S2 cells were prepared for imaging by gently transferring 1 mL of culture at ≈ 1×10^6^ cell/mL to a µ-Dish 35 mm imaging dish (Ibidi). The cells were allowed to adhere for 20-30 min prior to imaging by spinning disc confocal fluorescence microscopy (Nikon Ti2E with Yokogawa CSU X1 spinning disc). Images were take with a 60x oil immersion objective and 405 and 488 nm lasers were used to excite mTurquoise2 (His2A) and mNeonGreen (Orc1), respectively. Samples were scanned to identify mitotic cells with chromosomes aligned at the metaphase plate. The x,y coordinates of multiple mitotic cells were marked and the cellular dynamics of Orc1 assessed by time lapse imaging with a z-stack (8-14 µm thick section at 0.3 µm intervals) taken every 3 min with 300 ms exposure throughout mitosis.

FIJI was used for image processing and quantitation. For each image set, a maximum intensity projection was generated and a median filter was applied (pixel radius = 2) before the image was split into blue and green channels and background intensity subtracted. Regions of interest were generated for chromatin by auto thresholding on mTurquoise2 signal. The His2A and Orc1 chromosome intensity was measured within these regions for each time point. Similarly, cytosolic mNeonGreen-Orc1 intensity was measured for each time point. The fold change in chromosome intensity was calculated and normalized to the signal at metaphase. To determine the chromosome partitioning of Orc1, mNeonGreen-Orc1 signal on chromosomes was multiple by the chromosome area and divided by the sum of the same plus mNeonGreen-Orc1 cytosolic intensity times cytosolic area (as shown in **Fig 3A**). The average and standard deviation of Orc1 chromosome partitioning in metaphase and telophase was calculated for each construct in at least six different mitotic cells.

### Transfection and imaging of HeLa cells

The human Orc1^IDR^ coding sequence (residues 177-484) was inserted by LIC into vector 6D (QB3, Macrolab) which appends a GFP tag at the coding region’s C-terminus. HeLa cells stably expressing mCherry tagged human histone H2B (35) (provided by Dr. Bryan Gibson) were transfected with 6D-*Hs*Orc1^IDR^ using the jetPRIME transfection reagent (Polyplus) following the manufacturer’s protocol. Briefly, HeLa cells were cultured in Dulbecco’s Modified Eagle Media (DMEM) supplemented with 5 % FBS and maintained at 37°C, 5% CO_2_. A six-well plate was seeded with 2×10^6^ cells/well and incubated for 24 hr to achieve no more than 60-80% confluence. For the transfection, DNA was diluted in jetPRIME buffer, and then mixed with transfection reagent. The mixture was incubated at room temperature for 10 minutes before adding dropwise to cells and then gently mixing. The plate was incubated for 48 to 72 hours before imaging. To image, samples were transferred to a µ-Dish 35 mm imaging dish (Ibidi) and imaged by spinning disc confocal fluorescence microscopy (Nikon Ti2E equipped with Yokogawa CSU X1 spinning disc). The imaging set up was as described for S2 cells, with the exception that imaging was done with a 40x air objective and a 568 nm laser was used to excite mCherry. Images were processed as described for S2 cells.

### Cloning, expression, and purification of the ORC holocomplex and CDK/Cyc complexes

The ORC holocomplex (*Dm*ORC), the holocomplex lacking the Orc1 IDR (*Dm*ORC^Δ1IDR^), *Dm*CDK1/CycA, and *Dm*CDK2/CycE were expressed and purified from Sf9 cells. The coding region of *Dm*Orc1 was cloned into vector 438B (QB3 Macrolab) for expression with an N-terminal hexahistidine tag. *Dm*Orc2, *Dm*Orc3, and *Dm*Orc5 were cloned into vector 438A (no tag, QB3 Macrolab). Gibson cloning was used to clone *Dm*Orc4 into vector 438B replacing the hexahistidine coding sequence with a TEV-cleavable Maltose Binding Protein (MBP) tag. Orc1-5 were combined into a single multibac expression vector. *Dm*Orc6 was cloned into pFastBac. Plasmids for production of *Dm*ORC1-6^Δ1IDR^ were the same except that IDR residues 248-549 were deleted from 438B-*Dm*Orc1 and this was not combined in a multibac with Orc2-5. *Dm*CDK1 and CDK2 coding sequences were cloned into our custom 438-MBP vector for expression as an N-terminal TEV-cleavable MBP fusion. Cyclin subunits (CycA or CycE) were cloned into vector 438B. Bacmid DNA was generated from these vectors according to previously established methods (19). Subsuequntly, bacmid DNAs were transfected into Sf9 cells (Expression Systems) maintained in ESF921 using Cellfectin II following the manufacturer’s protocol (Fisher Scientific). P0 virus was harvested from the transfected cells and amplified twice prior to infection of Sf9 cells for protein expression. Cells were co-infected with multiple viruses for production of holocomplexes (*Dm*ORC, *Dm*ORC^Δ1IDR^, *Dm*CDK1/CycA, and *Dm*CDK2/CycE) and harvested after 2 days. Cell pellets were isolated by centrifugation and frozen at -80°C until protein purification.

*Dm*ORC was purified from 2 L of cultured cells and the cell pellet resuspended in 80 mL of Lysis Buffer (50 mM Tris pH 7.5, 300 mM KCl, 50 mM Imidazole, 10% glycerol, 200 µM PMSF, 1 mM BME, 1 µM benzonase and 1x cOmplete EDTA-free Protease Inhibitor Cocktail (Sigma-Aldrich)). Cells were lysed by sonicating on ice (sonicate program of 5 cycles of: 15 sec 100% power, 1 min rest) and the lysate centrifuged at 18,000 RPM for 1 hr at 4°C. The supernatant was collected and filtered using aPES 0.45 µm bottle-top filter unit (Nalgene Rapid-Flow, ThermoFisher) and then subject to an ammonium sulfate precipitation (975 mM ammonium sulfate) and incubated under gentle rotation for 30 min at 4°C. The lysate was further clarified by centrifugation at 18,000 RPM for 1 hr at 4°C. The sample was filtered again prior to Nickel purification. The filtrate was loaded onto a 5 mL HisTrap HP column (GE Healthcare), washed with 12 column volumes (CV) of Wash Buffer (50 mM Tris pH 7.5, 300 mM KCl, 50 mM Imidazole, 10% glycerol, 1 mM BME), and the protein eluted with a 6 CV linear gradient from 0-100% Elution Buffer (50 mM Tris pH 7.5, 300 mM KCl, 250 mM Imidazole, 10% glycerol, 1 mM BME). Amylose purification was used as a second affinity purification step. The sample was loaded onto a column packed with amylose resin and subsequently washed with 3 CV of Amylose Wash Buffer (50 mM Tris pH 7.5, 300 mM KCl, 10% glycerol, 1 mM BME) and eluted with 2 CV of Amylose Elution Buffer (50 mM Tris pH 7.5, 300 mM KCl, 10% glycerol, 1mM BME, 20 mM maltose). The His6-MBP tag was cleaved by addition of TEV (1/10 w/w) and a 12 hr incubation at 4°C. Finally, TEV digested protein was concentrated to < 2 mL using an Amicon Ultra-15 concentrator (Millipore) and then purified by size exclusion chromatography. Samples were loaded onto a HiPrep 16/60 S300 HR and run with 1.2 CV of Sizing Buffer (50 mM HEPES pH 7.5, 300 mM KGlutamate, 10 % glycerol, 1 mM BME). Peak fractions were assessed by SDS-PAGE and subsequently pooled, concentrated, aliquoted, and flash frozen in liquid nitrogen and stored in -80°C. The same purification process was applied to CDK/Cyc complexes except that the TEV digest and sizing exclusion steps were omitted from the workflow.

### Cloning, expression, and purification of Orc1 IDR constructs

*E. coli* BL21(DE3) cells were used to express and purify Orc1 IDRs, including *Dm*Orc1^IDR^ (*D. melanogaster* Orc1 residues 187-549), synthesized (Twist Biosciences) metazoan Orc1^IDR^s (*A. millepora* Orc1 residues 170-485, *B. plicatilis* Orc1 residues 1-158, *M. yessoensis* Orc1 residues 175-621, *D. gyrociliatus* Orc1 residues 169-412, *H. dujardini residues* 1-278, and *A. queenslandica* Orc1 residues 177-593) and *Dm*Orc1 IDR variants (Δ1 = deletion of residues 187-246, Δ2 = deletion of residues 247-306, Δ3 = deletion of residues 307-366, Δ4 = deletion of residues 367-426, Δ5 = deletion of residues 427-486, Δ6 = deletion of residues 487-549, and ΔBP = deletion of residues 520-533). Each Orc1 IDR coding sequence was cloned into QB3 Macrolab vector 1C for expression as a TEV-cleavable N-terminal His6-MBP fusion. *Dm*Orc1^IDR^ deletion constructs were generated by around-the-horn PCR based mutagenesis using 1C-*Dm*Orc1^IDR^ as a template. Each construct was transformed into BL21(DE3) cells (NEB). Subseuquently, cells were grown in liquid culture to an OD600 = 0.8 and expressed in overnight cultures at 20°C upon 1 mM IPTG induction. Cells were harvested by centriguation and the collected cell pellets were stored at −80°C until protein purification.

For protein purification, cells from 2 L of culture were resuspended in 80 mL of Lysis Buffer (20 mM Tris pH 7.5, 500 mM NaCl, 30 mM Imidazole, 10% glycerol, 200 µM PMSF, 1x cOmplete EDTA-free Protease Inhibitor Cocktail (Sigma-Aldrich), 1 mM BME and 0.1 mg/mL lysozyme) and sonicated on ice before centrifugation at 18,000 RPM for 1 hr at 4°C. The sample was applied to an aPES 0.45 µm bottle-top filter unit (Nalgene Rapid-Flow, ThermoFisher) and the filtrate loaded onto a 5 mL HisTrap HP column (GE Healthcare). The column was washed with 10 CV of Nickel Wash Buffer (20 mM Tris pH 7.5, 500 mM NaCl, 30 mM Imidazole, 10% glycerol, 200 µM PMSF, 1 mM BME) and the protein eluted with 7 CV of Nickel Elution Buffer (20 mM Tris pH 7.5, 150 mM NaCl, 500 mM Imidazole, 10% glycerol, 1 mM BME). The protein was then applied to a HiTrap Heparin HP column (GE Healthcare) which was then washed with 10 CV of Heparin Bind Buffer (20 mM Tris pH 7.5, 150 mM NaCl, 10% glycerol, 1 mM BME, 400 µM PMSF) and eluted with a 10 CV linear gradient of increasing salt from 150 mM to 1 M NaCl. Elution fractions were assessed by SDS-PAGE and the fractions containing non-proteolyzed protein were pooled and TEV digested overnight at 4°C. The free tag, uncleaved protein and TEV were subsequently removed by an ortho nickel affinity purification step and the sample concentrated to **<** 2 mL using a 10K Amicon Ultra-15 concentrator (Millipore). Finally, the sample was loaded on a HiPrep 16/60 Sephacryl S-300 HR column pre-equilibrated and run in Sizing Buffer (50 mM HEPES pH 7.5, 300 mM KGlutamate, 10 % glycerol, 1 mM BME). Peak fractions were assessed by SDS-PAGE before being pooled, concentrated, flash frozen in liquid nitrogen, and stored at −80°C.

### *In vitro* phosphorylation reactions

In general, the molar ratio of substrate to kinase (CDK/Cyc complex) in each phosphorylation reaction was 5:1. Specifically, 4 µM ORC was mixed with 0.8 µM of CDK/Cyc and 12 µM Orc1^IDR^ was mixed with 2.4 µM CDK/Cyc in the presence or absence of 1 mM ATP in phosphorylation reaction buffer (50 mM HEPES pH 7.5, 300 mM KGlutamate, 10% glycerol, 5 mM MgOAc, 1 mM BME). The reaction mixtures were incubated for 30 mins at 25°C. Phosphorylation was assessed by SDS-PAGE with Coomassie staining where a shift in the band of the phosphorylated protein compared to the unphosphorylated control confirmed the success of the reaction.

We also performed a large scale preparation of phosphorylated *Dm*Orc1^IDR^. In this case, the size exclusion chromatography peak fractions containing purified *Dm*Orc1^IDR^ were pooled, concentrated, and then subject to CDK2/CycE-dependent phosphorylation. The phosphorylation reaction mixture contained a 5:1 molar ratio of *Dm*Orc1^IDR^ to CDK2/CycE in addition to 4 mM ATP in reaction buffer (50 mM HEPES pH 7.5, 300 mM KGlutamate, 10% glycerol, 5 mM MgOAc, 1 mM BME). The mixture was incubated for 30 mins at room temperature (25°C). Afterwards, the reaction mixture was further supplemented with 2 mM ATP and CDK2/CycE at a molar ratio of 20:1 and then incubated for an additional 30 min to drive the phosphorylation reaction to completion. The reaction mixture was then subjected to size exclusion chromatography in order to remove CDK2/CycE and ATP. Peak fractions containing phosphorylated *Dm*Orc1^IDR^ (p*Dm*Orc1^IDR^) were assessed by SDS-PAGE and then pooled, concentrated, flash frozen in liquid nitrogen, and stored at -80°C.

### Analysis of DNA binding by electrophoretic mobility shift assays (EMSAs)

Serial dilutions of Orc1 IDR variants were prepared in assay buffer (50 mM HEPES pH 7.5, 150 mM KGlutamate, 10% glycerol, 1 mM BME) containing 2 nM Cy5-dsDNA (5’-GAAGCTAGACTTAGGTGTCATATTGAACCTACTATGCCGAACTAGTTACGAGCTATAACC-3’). The mixtures were incubated at room temperature for 20 min and then run on a 1% agarose gel at 100 V for 30 min. The gel was imaged on a BioRad ChemiDocMP using imaging settings appropriate for the excitiation and emission spectra of the fluorescent label (Cy5 or FITC). The acquired images were analyzed by quantitating the loss of free DNA and calculating the fraction bound for each protein concentration. These data were plotted in Graphpad Prism and fit with a Hill Equation to determine the dissociation constant. The data from three independent experiments were used to calculate mean and standard deviation, which are reported.

### Analysis of DNA binding by fluorescence polarization

Fluorescence anisotropy was performed using a FITC labeled dsDNA (5’-GAAGCTAGACTTAGGTGTCATATTGAACCTACTATGCCGAACTAGTTACGAGCTATAACC-3’). Reactions invariably contained 2 nM FITC-dsDNA but varying levels of KGlut (150 or 300 mM) and ATP (0 mM or 1 mM). The remaining buffer components were always the same: 50 mM HEPES pH 7.5, 10% glycerol, 5mM MgOAc, 1 mM BME. Concentrations of ORC and Orc1^IDR^ are as indicated in figure legends. DNA/protein mixtures were incubated for 30 minutes at room temperature before being added to 384-well black bottom multi-well plate and transferred to a CLARIOstar BMG LABTECH plate reader for fluorescence polarization measurement and calucation of fluorescence polarization. All data were background corrected for a buffer blank before calculating polarization. At least 3 biological replicates, each with technical duplicates, were performed and the mean and standard deviation of three independent experiments were plotted with respect to each protein concentration. The data were analyzed by Graphpad Prism by fitting a Hill Equation to the data to calculate dissociation constants.

Time-resolved fluorescence polarization DNA-binding assays were carried out for *Dm*Orc1^IDR^ at 150 mM KGlut in the presence of CDK2/CycE. The reaction mixture contained 5 µM *Dm*Orc1^IDR^, 50 nM Cdk2/CycE, 5 mM MgOAc, and 2 nM of FITC-dsDNA. The phosphorylation reaction was initiated by adding 1 mM ATP, gentle mixing, and then immediately and repeatedly taking fluorescence polarization readings. The control reaction was set up the same way but with the exclusion of ATP. Fluorescence polarization measurements were taken every 2.5 min for approximately 2 hours and measurements were plotted as function of time.

### Mass spectrometry analysis of CDK/Cyc-dependent phosphorylation

Intact protein samples (50 µL 0.3 mg/mL) were analyzed by LC/MS, using a Sciex X500B QTOF mass spectrometer coupled to an Agilent 1290 Infinity II HPLC. Samples were injected onto a POROS R1 reverse-phase column (2.1 × 30 mm, 20 µm particle size, 4000 Å pore size) and desalted. The mobile phase flow rate was 300 uL/min and the gradient was as follows: 0-3 min: 0% B, 3-4 min: 0-15% B, 4-16 min: 15-55% B, 16-16.1 min: 55-80% B, 16.1-18 min: 80% B. The column was then re-equilibrated at initial conditions prior to the subsequent injection. Buffer A contained 0.1% formic acid in water and buffer B contained 0.1% formic acid in acetonitrile. The mass spectrometer was controlled by Sciex OS v.3.0 using the following settings: Ion source gas 1 30 psi, ion source gas 2 30 psi, curtain gas 35, CAD gas 7, temperature 300 °C, spray voltage 5500 V, declustering potential 135 V, collision energy 10 V. Data was acquired from 400-2000 Da with a 0.5 s accumulation time and 4 time bins summed. The acquired mass spectra for the proteins of interest were deconvoluted using BioPharmaView v. 3.0.1 software (Sciex) in order to obtain the molecular weights. The peak threshold was set to ≥ 5%, reconstruction processing was set to 20 iterations with a signal-to-noise threshold of ≥ 20 and a resolution of 2500.

For identification of phospho-sites, samples were digested overnight with trypsin (Pierce) following reduction and alkylation with DTT and iodoacetamide (Sigma–Aldrich). The samples then underwent solid-phase extraction cleanup with an Oasis HLB plate (Waters) and the resulting samples were injected onto a QExactive HF mass spectrometer coupled to an Ultimate 3000 RSLC-Nano liquid chromatography system. Samples were injected onto a 75 um i.d., 15-cm long EasySpray column (Thermo) and eluted with a gradient from 0-28% buffer B over 90 min with a flow rate of 250 nL/min. Buffer A contained 2% (v/v) ACN and 0.1% formic acid in water, and buffer B contained 80% (v/v) ACN, 10% (v/v) trifluoroethanol, and 0.1% formic acid in water. The mass spectrometer operated in positive ion mode with a source voltage of 2.5 kV and an ion transfer tube temperature of 275 °C. MS scans were acquired at 120,000 resolution in the Orbitrap and up to 20 MS/MS spectra were obtained for each full spectrum acquired using higher-energy collisional dissociation (HCD) for ions with charges 2-8. Dynamic exclusion was set for 20 s after an ion was selected for fragmentation.

Raw MS data files were analyzed using Proteome Discoverer v2.4 SP1 (Thermo), with peptide identification performed using Sequest HT searching against the *Drosophila melanogaster* protein database from UniProt along with the sequence of Orc1^IDR^. Fragment and precursor tolerances of 10 ppm and 0.02 Da were specified, and three missed cleavages were allowed. Carbamidomethylation of Cys was set as a fixed modification, with oxidation of Met and phosphorylation of Ser, Thr, and Tyr set as variable modifications. Phosphorylated sites were localized using the IMP-ptmRS node within Proteome Discoverer. The false-discovery rate (FDR) cutoff was 1% for all peptides.

### Bioninformatic analysis of Orc1 IDR orthologs

Sequence alignments were done with Geneious software (Dotmatics). Pairwise comparisons of the compositional differences between metazoan Orc1 orthologs were performed using a modified version of the chi-square test of homogeneity that quantifies differences in the fractional content of amino acids (29). Instead of using the test statistic to confirm or reject a null hypothesis, the score is normalized so that compositionally identical sequences receive a score of zero while sequences with no residues in common receive a score of one. Applied intra-sequentially, the chi-score analysis was used to identify regions of distinct amino acid composition within the *D. melanogaster* Orc1-IDR. This was done by using the pairwise chi-scores between subsequences to predict the positions of module boundaries, which were then optimized and validated statistically. Boundaries were iteratively removed until only those with z-scores corresponding to a confidence level of 95% or higher remained. The application of the chi-square method to compare sequence composition between proteins and local bias within a contiguous region of disorder is described in detail in a recent publication (29).

## ACKNOWLEDGEMENTS

We thank members of the Parker lab and the UTSW Condensate Club, including Rosen, Sabari, and Woodruff labs, for helpful discussion and advice. We also thank Andrew Lemoff of the UTSW Proteomics Core for guidance in assessing ORC phosphorylation by mass spectrometry. M.W.P. is the Cecil H. and Ida Green Endowed Scholar in Biomedical Computational Science. This work was supported by The Welch Foundation (I-2074-20210327, to M.W.P.) and by the Cancer Prevention and Research Institute of Texas (CPRIT, RR200070, to M.W.P.).

## FIGURES, FIGURE TITLES AND LEGENDS

**S1 Fig:**
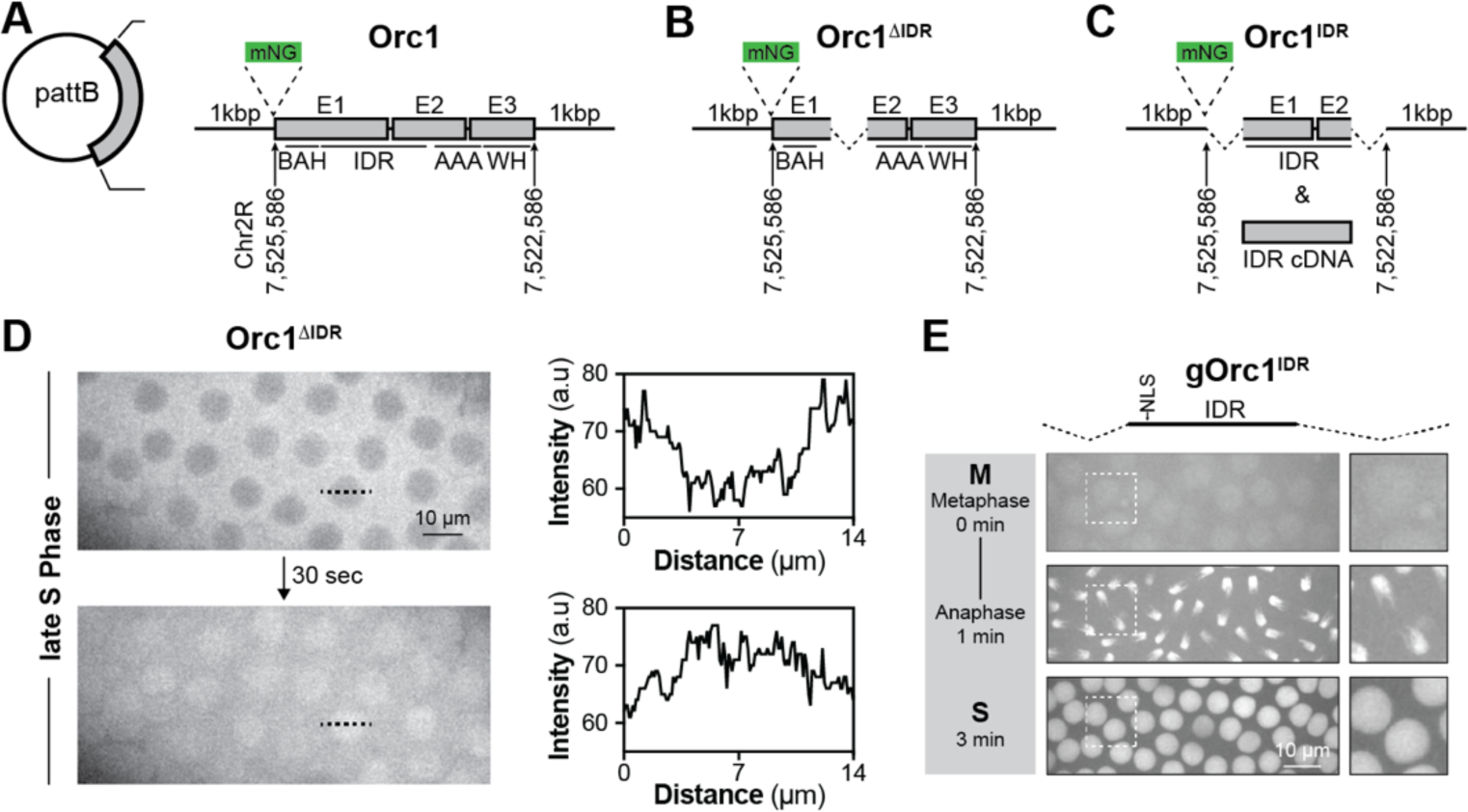
The fly Orc1 IDR is necessary and sufficient for chromatin recruitment in vivo. A-C) Graphical representation of Orc1 transgenes generated in vector pattB, including full-length Orc1 (Orc1, A), an IDR deletion construct (Orc1^ΔIDR^, B), and the IDR alone (Orc1^IDR^, C). D) Imaging in embryos in late S-phase reveals a dramatic change in Orc1^ΔIDR^ nuclear localization. Included are line intensity profiles of mNeonGreen-Orc1^ΔIDR^ signal. E) Chromatin localization of an Orc1^IDR^ transgene produced from genomic DNA.

**S2 Fig:**
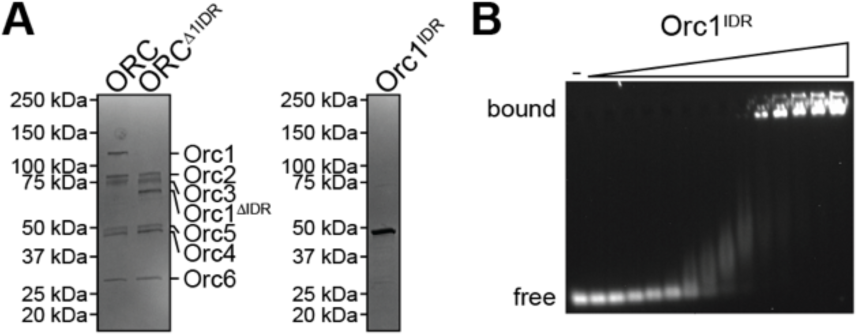
The Orc1 IDR facilitates ATP-independent DNA binding in vitro. A) SDS-PAGE of purified DmORC holocomplex (ORC) and an Orc1 IDR deletion construct (ORC^Δ1IDR^). Second gel is SDS-PAGE of purified DmOrc1^IDR^. B) EMSA analysis of DmOrc1^IDR^ DNA-binding. Each lane represents a 2-fold dilution of Orc1 (0.6 – 5,000 nM) which was combined with 2 nM Cy5-dsDNA in Assay Buffer (50 mM HEPES pH 7.5, 150 mM KGlutamate, 10% glycerol, 1 mM BME).

**S3 Fig:**
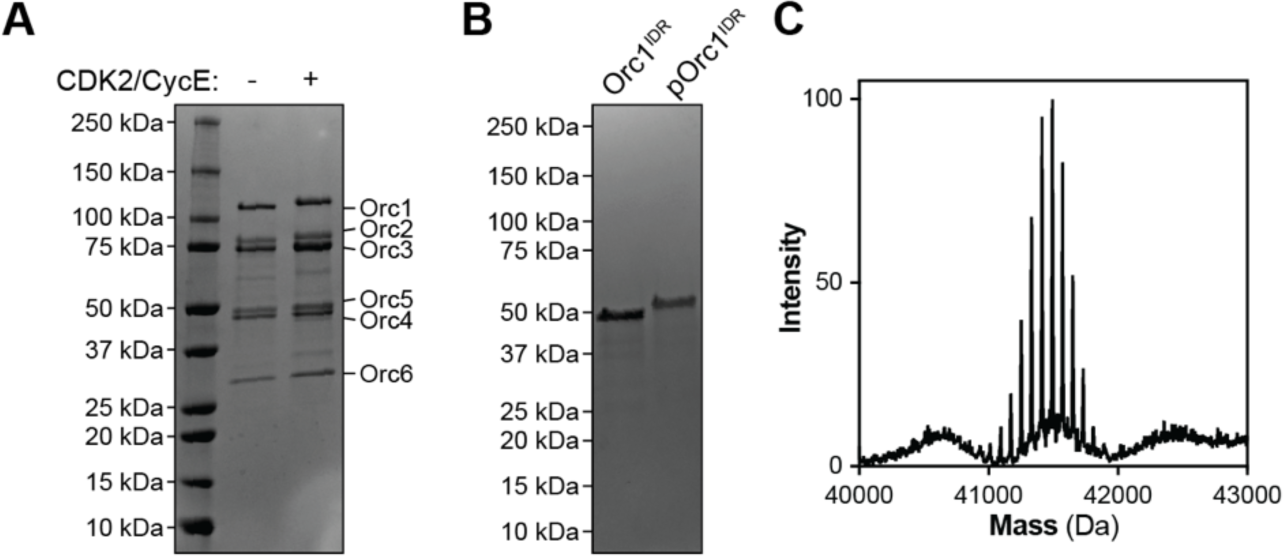
Phosphorylation of the Orc1 IDR regulates DNA and chromatin binding. A) SDS-PAGE analysis of non-phosphorylated (“-”) and CDK2/CycE phosphorylated ORC (“+”). B) SDS-PAGE analysis of purified Orc1^IDR^ and CDK2/CycE phosphorylated Orc1^IDR^ (pOrc1^IDR^). C) Intact mass spectrometry reveals that pOrc1^IDR^ has been phosphorylated approximately 15 times.

**S4 Fig:**
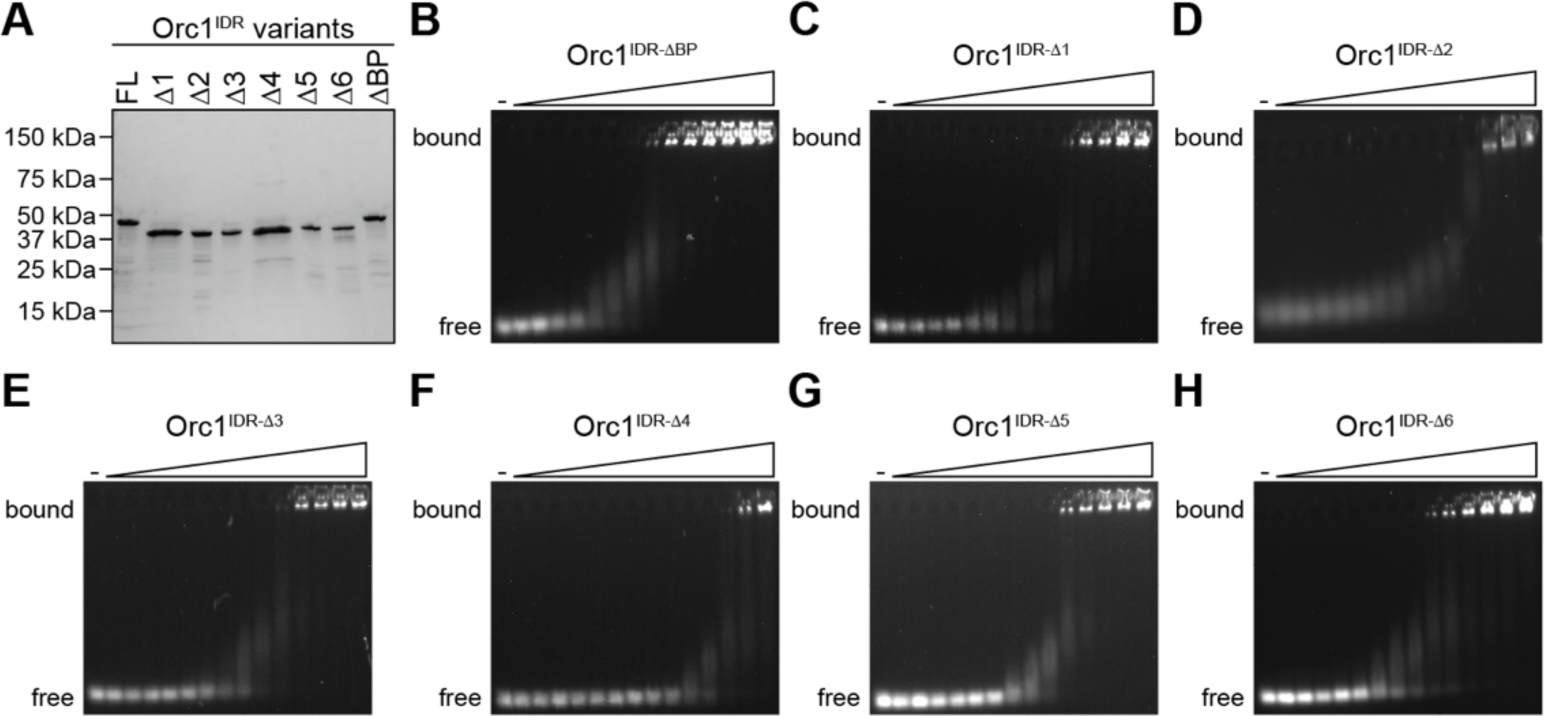
The Orc1 disordered region possesses multiple redundant DNA binding motifs. A) SDS-PAGE analysis of purified Drosophila Orc1^IDR^ and the listed deletion constructs. See MATERIALS AND METHODS for indices of deleted amino acids. B-H) EMSA analysis of Orc1^IDR^ deletion construct binding to Cy5-dsDNA.

**S6 Fig:**
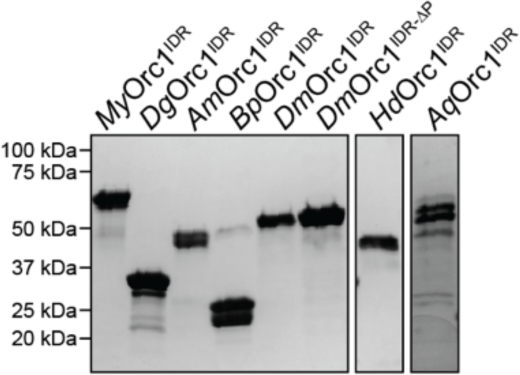
The Orc1 IDR is functionally conserved across metazoans. SDS-PAGE of purified metazoan Orc1 IDRs (Mollusc (My), Annelid (Dg), Cnidaria (Am), Rotifer (Bp), Arthropod (Dm and DmOrc1^IDR-ΔP^, Tardigrade (Hd), and Porifera (Aq)).

